# Mycobacteria form viable cell wall-deficient cells that are undetectable by conventional diagnostics

**DOI:** 10.1101/2022.11.16.516772

**Authors:** Noortje Dannenberg, Victor J. Carrion Bravo, Tom Weijers, Herman P. Spaink, Tom H. M. Ottenhoff, Ariane Briegel, Dennis Claessen

**Affiliations:** Institute of Biology, Leiden University, Sylviusweg 72, 2333 BE Leiden, The Netherlands; Departamento de Microbiología, Instituto de Hortofruticultura Subtropical y Mediterránea ‘La Mayora’, Universidad de Málaga-Consejo Superior de Investigaciones Científicas (IHSM-UMA-CSIC), Universidad de Málaga, Málaga, Spain; Department of Microbial Ecology, Netherlands Institute of Ecology (NIOO-KNAW), Wageningen, the Netherlands; Infectious Diseases, Leiden University Medical Center, Albinusdreef 2, 2333 ZA Leiden, The Netherlands; Netherlands Centre for Electron Nanoscopy (NeCEN), Leiden University, Einsteinweg 55, 2333 CC Leiden, The Netherlands

**Keywords:** Mycobacteria, cell wall-deficiency, viability, antibiotics, osmolality, diagnostics

## Abstract

The cell wall is a unifying trait in bacteria and provides protection against environmental insults. Therefore, the wall is considered essential for most bacteria. Despite this critical role, many bacteria can transiently shed their cell wall and recent observations suggest a link of such wall-deficient cells to chronic infections. Whether shedding the cell wall also occurs in mycobacteria has not been established unambiguously. Here we provide compelling evidence that a wide range of mycobacterial species, including clinical and non-clinical isolates, form viable cell wall-deficient cells in response to environmental stressors. Using cryo-transmission electron micrography we show that the complex multi-layered wall is largely lost in such cells. Notably, we show that their formation in *Mycobacterium marinum* and BCG vaccine strains of *Mycobacterium bovis* is stimulated by exposure to cell wall-targeting antibiotics. Given that these wall-deficient mycobacteria are undetectable using conventional diagnostic methods, such cells have likely been overlooked in clinical settings. Altogether, these results indicate that mycobacteria can readily switch between a walled and wall-deficient lifestyle, which provides a plausible explanation for enabling persistence of infections caused by members of this genus.

## INTRODUCTION

Mycobacteria are the causative agents of some of the most serious human infectious diseases, including tuberculosis (TB), leprosy and a range of other infections (WHO, 2020; WHO, 2021). Upon transmission, these pathogens cause progressive infections that need to be treated with elaborate antibiotic regimes. However, such treatments may stimulate persistence in the host and promote the development of antibiotic resistance (Simmons *et al.,* 2018, Johansen *et al.,* 2020, Saxena *et al*., 2021). Tackling persistence is challenging because these cells are difficult to cultivate and phenotypically heterogeneous. Therefore, many fundamental questions regarding their physiological state and mechanism of survival remain unanswered (Barry *et al*., 2009; Dhar and McKinney, 2007; Manina *et al*., 2014).

Despite being viewed with scepticism, some studies have indicated that mycobacterial persister cells could be cell wall-deficient (CWD), although these studies have been difficult to replicate due to extreme long incubation times of the *in vitro* persistence models (Shleeva *et al*., 2011; Velayati *et al*., 2016). The cell wall is a unifying trait in the bacterial domain and considered essential for most bacteria (Vollmer *et al*., 2008). As such, the enzymes involved in cell wall synthesis are among the prime targets of effective antibiotics, which typically act by preventing normal cell wall assembly and thereby causing cell death. Considering the importance of the cell wall, it is surprising that under specific stressful conditions a range of bacteria can transiently escape from their enclosing cell wall to adopt a viable but CWD lifestyle (Allan *et al*., 2009; Claessen and van Wezel, 2014; Errington, 2013, 2017; Errington *et al*., 2016; Kawai *et al*., 2015; Klieneberger, 1935; Mercier *et al*., 2014; Ramijan *et al*., 2018; Claessen and Errington, 2019, Lazenby *et al.,* 2022). These cells are competent with regard to DNA uptake (Woo *et al.,* 2003, Kapteijn *et al.,* 2022) and can revert to their walled state when the stressful conditions have diminished. Furthermore, these cells can oftentimes proliferate without their cell wall, as so-called “L-forms”, under specific growth conditions or through the acquirement of beneficial mutations. Such mutations are known to reduce susceptibility to reactive oxygen species (ROS) and to increase membrane fluidity. Importantly, the loss of the cell wall provides a mechanism via which bacteria are protected from cell wall-targeting antibiotics (Kawai *et al*., 2018). Besides, as the most important antigens of bacteria are located on the cell surface, loss of the cell wall could potentially facilitate immune evasion and contribute further to persistence in the host (Källenius *et al*., 2016).

Recent observations suggest a link between cell wall-deficiency and chronic infections, based on the isolation of *Escherichia coli* CWD cells from patients suffering from recurrent urinary tract infections (Mickiewicz *et al*., 2019). This raises the question whether CWD may also play a role in other infectious diseases, such as tuberculosis. In this study, we show that multiple mycobacterial species can naturally form viable CWD cells in response to environmental stressors. Confocal microscopy and cryo-transmission electron microscopy confirmed that these cells contain DNA but lack major cell wall structures. Importantly, in *Mycolicibacterium smegmatis,* mycobacterial endophyte isolates (Carrion *et al*., 2019), clinical isolates of the *Mycobacterium avium* complex, *Mycobacterium marinum* strains and *Mycobacterium bovis* BCG vaccine strains formation of these cells is stimulated by the presence of cell wall-targeting agents, including isoniazid, D-cycloserine and vancomycin, of which the former two are used clinically to treat infections. Furthermore, we show that conventional diagnostics fail to detect CWD cells, suggesting that such cells may have been over-looked in clinical diagnostic settings. Altogether, our work provides important insights in morphological and phenotypic plasticity in mycobacteria, which in the future may be exploited to target these devastating pathogens.

## RESULTS

### Mycobacteria produce wall-deficient cells under hyperosmotic stress conditions

To investigate the ability of Mycobacteria to generate CWD cells, we grew the fast-growing model organism *Mycolicibacterium smegmatis* mc^2^155 in media containing high levels of osmolytes, which was previously used for the formation of CWD cells in other actinobacteria (Ramijan *et al.,* 2018). Interestingly, growth of *M. smegmatis* on solid L-Phase Media Agar (LPMA) resulted in a change in colony morphology. Whereas on standard Middlebrook 7H10 medium colonies appeared rough and white, those on LPMA were small and transparent (Fig. 1A). Further investigation by light microscopy revealed spherical, vesicular structures between the elongated mycobacterial cells on LPMA, which were not found when using 7H10 medium. Similar vesicles were also identified when the strain was grown in liquid L-Phase Broth (LPB) and these vesicles contained nucleic acids as revealed by labelling with SYTO 9 (Fig. 1B). By using a constitutively expressing mCherry strain, these vesicles were found to contain fluorescent cytosol (Fig. 1C, S1D), perhaps suggesting that they are metabolically active cells. Indeed, these cells increased in size over time, from an average of 1.03 ± 0.54 µm (n = 561) to 2.15 ± 0.97 µm (n = 684) after 1 day and 10 days, respectively. Some cells even reached a size of more than 5 µm, coinciding with an increased complexity of the cellular content (Fig. 1D, S1).

**Figure 1.**
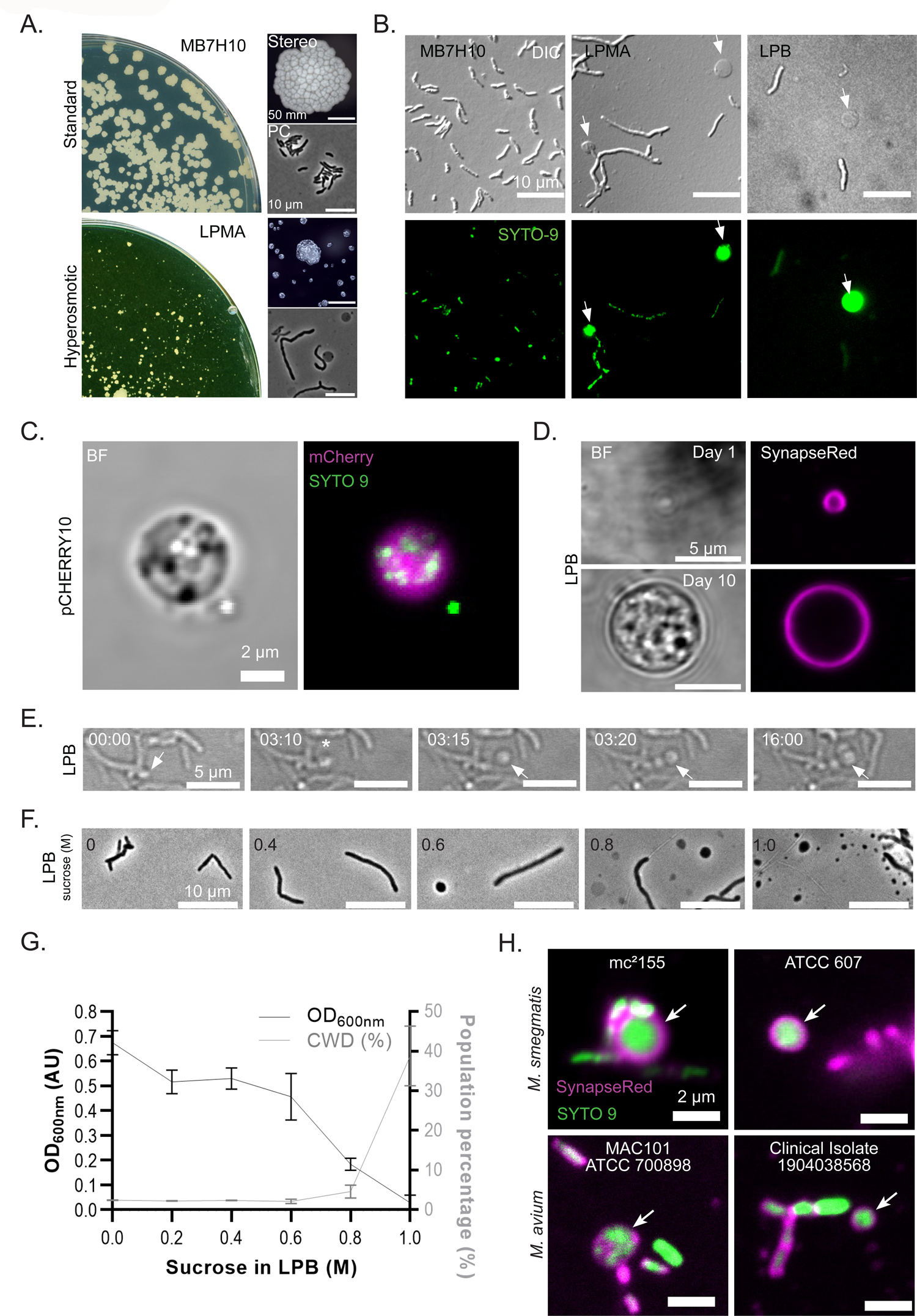
Hyperosmotic stress causes cell wall-deficiency in mycobacteria. A) Plate scans, stereo images and phase contrast micrographs of *M. smegmatis* mc^2^155 colonies grown for 3 days on standard cultivation medium MB7H10 or for 6 days on hyperosmotic LPMA medium. Scale bar represents 50 mm and 10 µm. B) DIC and fluorescent micrographs of *M. smegmatis* mc^2^155 grown for 3 days on MB7H10 and LPMA plates and in liquid LPB medium. Nucleic acids were labeled with SYTO 9 for 15 min. Scale bars represent 10 µm. C) Confocal image of a nucleic acid-containing cell of *M. smegmatis* mc^2^155 fluorescently labeled with pCHERRY10 (Caroll *et al.,* 2010) after 10 days of growth in LPB medium. Nucleic acids were labeled with SYTO 9 for 15 min. Scale bar represents 2 µm. D) Confocal images of *M. smegmatis* mc^2^155 cells with different diameters, 1.8 µm and 6.8 µm, respectively, formed in LPB medium. Membranes were labeled with SynapseRed for 15 min. Scale bars represent 5 µm. E) Stills of *M. smegmatis* mc^2^155 showing extrusion of CWD cells in LPB medium. Scale bars represent 5 µm. F) Phase contrast micrographs of *M. smegmatis* mc^2^155::pCHERRY10 grown in ascending concentrations of sucrose for 1 day. Scale bars represent 10 µm. G) Correlation between optical density and the number of CWD cells of *M. smegmatis* mc^2^155::pCHERRY10 in LPB supplemented with ascending concentrations of sucrose. The number of CWD cells is represented as the percentage of all cells in the cultures, which was quantified using flow cytometry (See Figs. S1E and S2). Optical density measurements (OD_600nm_) was were performed over time as shown in Supplemental Figure S1F. H) Confocal images of mycobacterial strains grown in LPB supplemented with 1.0 M sucrose. Images were made after 2 days (*M. smegmatis* strains) or 2 weeks (*Mycobacterium avium* complex (MAC) strains). Nucleic acids and cell membrane were labeled with SYTO 9 and SynapseRed, respectively, for 15 min. Scale bars represent 2 µm.

To gain insight into the origin of the spherical cells in the context of the rod-shaped cells, we performed live imaging (Fig. 1E, Video S1). These experiments revealed that these cells were extruded from the tips of the polar growing cells. In addition, their abundance positively correlated with an increased sucrose concentration in the medium (Fig. 1F, S1E, S2), while inversely correlating to growth of the strain (Fig. 1G, S1F). Importantly, formation of these cells is not restricted to *M. smegmatis* mc^2^155. We also noticed that the parental strain isolated from human samples, *Mycolicibacterium smegmatis* ATCC607, forms such spherical cells in the presence of high sucrose concentrations (Fig. 1H). Likewise, such cells were found in clinical, veterinary and lab strains of *Mycobacterium avium* Complex (MAC), as well as endophytic *Mycobacterium* isolates from sugar beet (Fig. 1H, S3A). Interestingly, the veterinary specimen *Mycobacterium avium* subsp. *Hominissuis* 20-935 /2, isolated from pig liver, produced an excess amount of membrane structures, but the majority did not contain nucleic acids (Fig. S3B).

**Figure 2.**
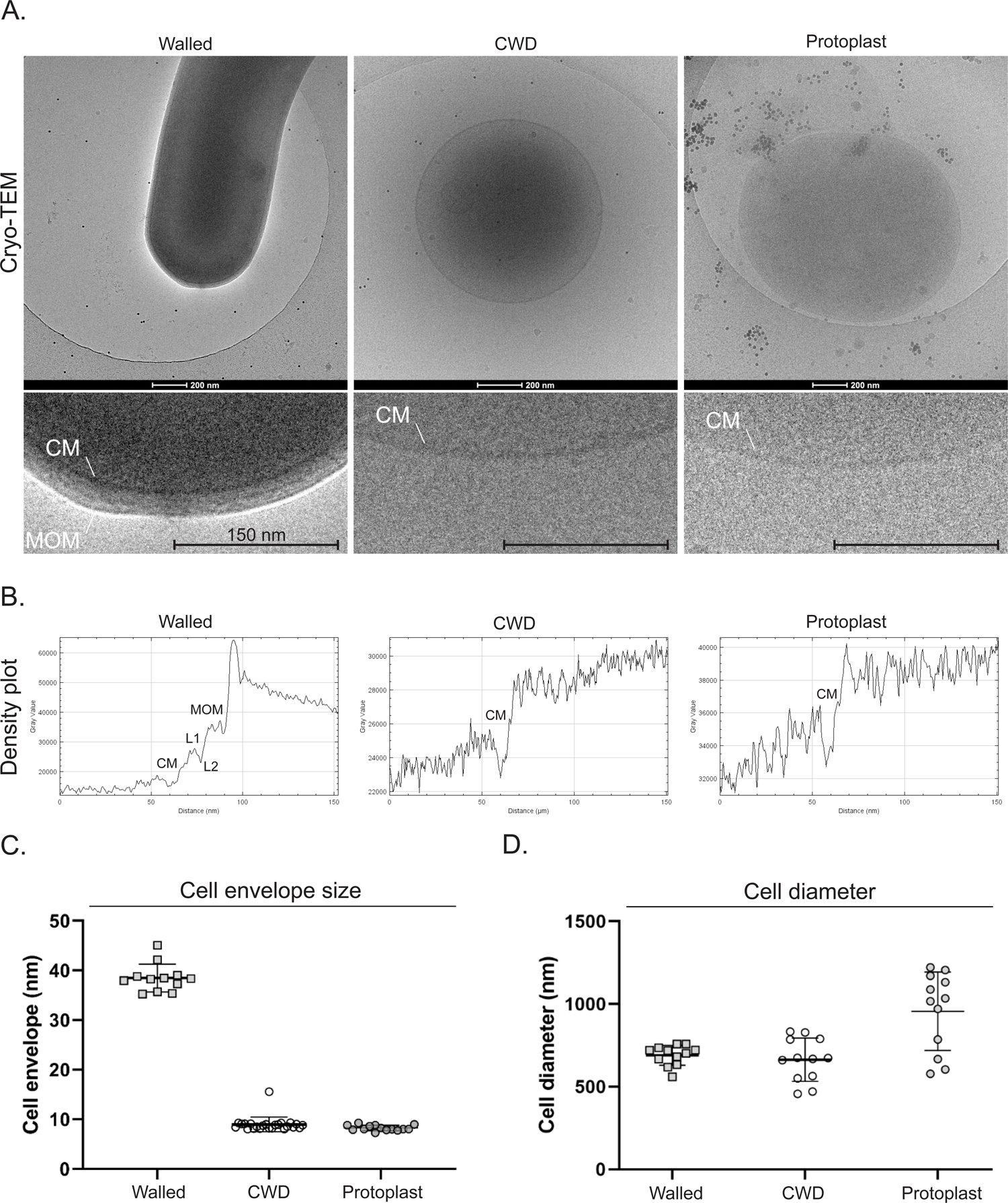
Wall-deficient cells of *M. smegmatis* lack major cell wall components. **A)** Cryo-Transmission Electron Microscopy (TEM) images of walled cells, CWD cells or protoplasts of *M. smegmatis* mc^2^155. Walled cells were grown in MB7H9 liquid medium and blotted directly from culture. CWD cells were enriched after 8 days of growth in LPB 1.0 M sucrose. Protoplasts were generated by lysozyme treatment in P-buffer through an adjusted mycobacterial protoplast protocol (see Material and Methods). Scale bars represent 200 nm. The lower panels represent zoomed in areas and scale bars correspond to 150 nm. CM, cytoplasmic membrane; MOM, mycobacterial outer membrane (Sani *et al.,* 2010). **B)** Gray scale density plot measurements from zoomed-in Cryo-TEM micrographs across the cell envelope in right-angle (90°). CM, cytoplasmic membrane; L1 and L2, domain-rich periplasmic layers; MOM, mycobacterial outer membrane (Hoffmann *et al.,* 2008). **C)** Cell envelope measurements based on density plots of selected cells (n = 6). Measurements were performed in duplicate at different locations per cell. The envelope diameter of CWD cells and protoplasts were measured in duplicate per cell (in 90° angle) and the walled cells were measured at the tip and at the side wall in relative distance from the tip curvature. **D)** Cell diameter measurements of selected cells (n=6). The diameter was measured in duplicate per cell, in perpendicular for spherical cells (CWD cells and protoplast) and perpendicular to the longitudinal axis of the rod shaped cell (side to side walls), with an minimum of 50 nm distance (Walled).

To characterize the spherical cells of *M. smegmatis* in more detail we successfully enriched them through filtration and concentration steps (Fig. S4) and analysed them in their native state. Cryo-transmission electron micrography (TEM) revealed that these cells lack major mycobacterial cell wall components, similarly to protoplasts, in which the cell wall is artificially removed through lysozyme treatment (Fig. 2A, S5). Comparable to protoplasts, the CWD cells are enveloped by the lipid bilayer, which is around 10 nm as seen in the grey value density plot, compared to the 45 nm cell envelope of walled cells (Fig. 2B, 2C). In some CWD cells, patches of cell wall material are evident exterior to the cell membrane (Fig. S5D). The diameter of these CWD cells range from 450 – 830 nm, which is similar to protoplasts that are 550-1200 nm in size (Fig. 2D). However, these CWD cells are considerably larger than extracellular vesicles, which typically range from 60 – 300 nm (Prados-Rosales *et al*., 2011; 2014; Gupta and Rodrigues, 2018). Furthermore, unlike extracellular vesicles, the content of CWD cells is dark-phased, suggesting dense packaging of macro-molecules. Altogether, these results demonstrate that formation of CWD cells in mycobacteria is a widespread mycobacterial response to hyperosmotic stress conditions.

### Mycobacterial wall-deficient cells are viable

To analyse if the CWD cells are viable and able to revert to their walled state, we attempted to separate them from walled cells using Fluorescent Activated Cell Sorting (FACS). However, the sensitive nature of the CWD cells complicated cell sorting, as these structures do not withstand the lack of osmo-protectants in the sheet fluid (Data not shown). Therefore, we indirectly tested their viability by removal of the CWD cells through the addition of the non-ionic detergent Triton X-100 (TX-100), a routinely used agent to solubilize membranes (Mattei *et al*. 2017). This treatment results in the formation of pores in the lipid bilayer membrane, causing explosive lysis, while cell-wall containing cells are protected. If the CWD cells are indeed viable, then treating such cells with detergent should result in a measurable drop of colony forming units (CFUs).

In accordance with this hypothesis, exposure to TX-100 removed all mCherry labelled CWD cells, while walled cells appeared unaffected (Fig. 3A). Real-time imaging indeed revealed the removal of the fluorescent CWD cells over time (Fig. 3B, Video S2), which was further substantiated by flow cytometric analysis (Fig. 3C, CWD cells: Q3; Pearson Chi-Square test, df = 2; χ^2^ =25.15, *p-*value = 3.45e^-06^; Table S3). Importantly, exposure to TX-100 reduced the formation of colony forming units 60-fold (Paired student t-test, tails 2: *p-*value= 0.012; Table S4), similarly to treatment of protoplasts with this detergent. By contrast, no decline in viability was observed when walled cells were exposed to TX-100. Taken together, these data demonstrate that a substantial fraction of CWD cells contributes to bacterial viability and replication competence as determined by CFU counts.

**Figure 3.**
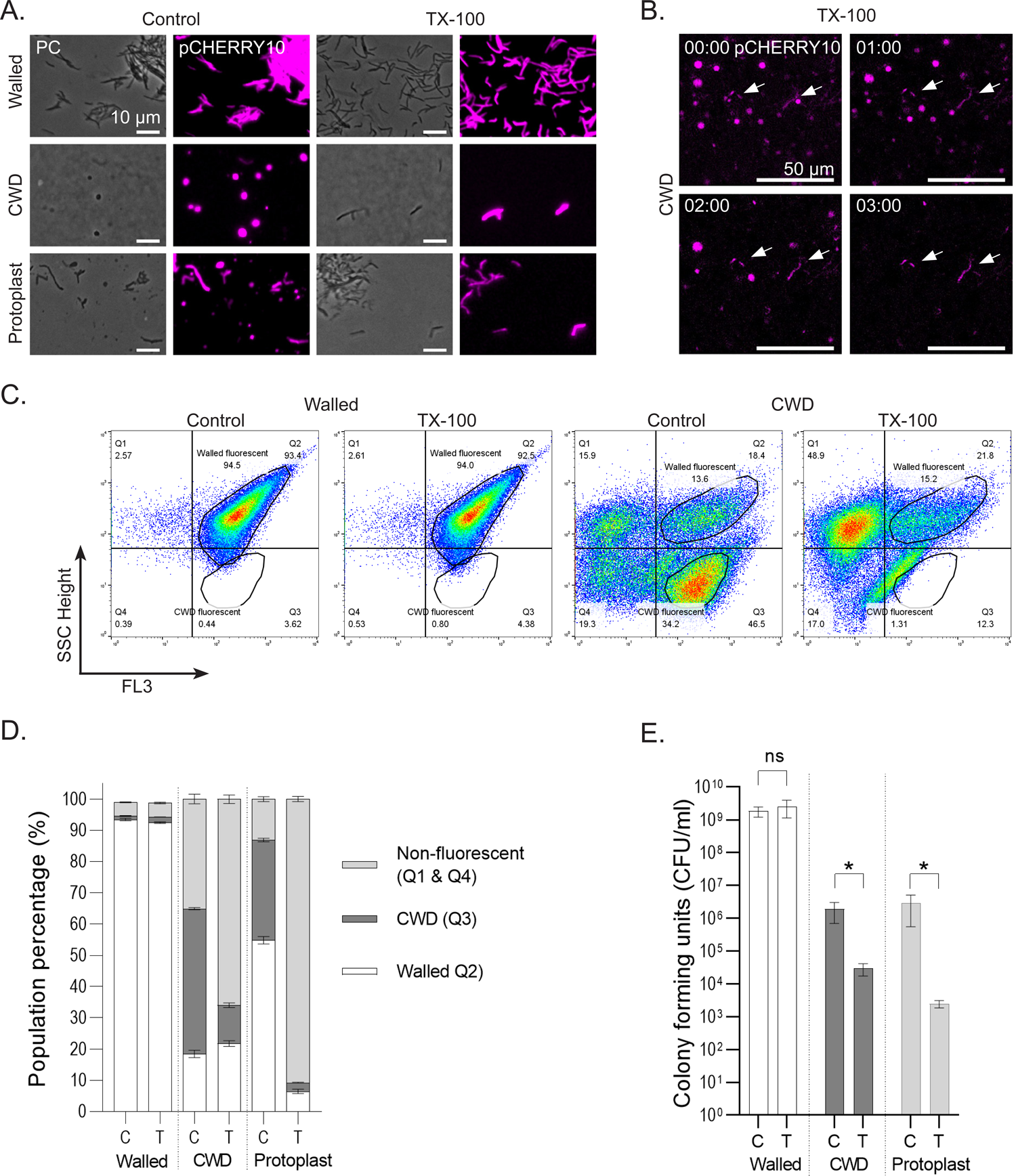
Wall-deficient cells of *M. smegmatis* are viable. **A)** Phase contrast and fluorescent images of *M. smegmatis* mc^2^155::pCHERRY10 walled cells, CWD cells and protoplasts after exposure to detergent 0.1% Triton X-100 (TX-100) for 15 min. As a control MQ was added to the cells. Scale bars represent 10 µm. **B)** Time-lapse images of *M. smegmatis* mc^2^155::pCHERRY10 exposed to 0.1 % TX-100. Please note that walled cells, indicated with arrows, do not disappear over time in contrast to CWD cells. Scale bars represent 50 µm. Time is indicated in hours. **C)** Flow cytometry plots of M. smegmatis mc^2^155::pCHERRY10 walled and CWD cells in response water (control) or t Triton X-100 exposure (TX-100) in triplicates. Plots were divided in four quadrants representing Highly complex, non-fluorescent particles (Q1), Highly complex fluorescent particles (Q2, Walled cells), Less complex, fluorescent particles (Q3, CWD cells), and Less complex, non-fluorescent particles (Q4). Please note that the CWD cells disappear when treated with TX-100 (right panels). **D)** Quantification of flow cytometry data. White bars represent walled cells, while light grey and dark grey represent non-fluorescent particles and CWD cells, respectively. Please note that the addition of TX-100 (T), in contrast to the water control (C), dramatically reduces the number of CWD cells and protoplasts, while having no effect on the walled cells. All measurements were performed in triplicate (see Fig. S6). **E)** Quantification of Colony Forming Units (CFU) counts of walled cells, CWD cells and protoplasts after the exposure to TX-100 (T) or water (C). Colonies were counted after 3 days. All measurements were performed in triplicate.

### Antibiotics induce formation of CWD cells in lower sucrose media

Cell wall-targeting antibiotics are a mainstay in treatment strategies of mycobacterial infections. To analyse the effect of such antibiotics on formation of CWD cells, we treated the cell cultures with the mycolic acid-targeting compound isoniazid (INH) and the peptidoglycan targeting compounds D-cycloserine (DC) and vancomycin (Van). Interestingly, exposure to these three antibiotics stimulated the formation of CWD cells in *M. smegmatis* mc^2^155 (Fig. 4A). Notably, in the presence of these antibiotics CWD cells became even evident in media with lower levels of sucrose (Fig. 4A, 4B, S7, S8). However, we never detected CWD cells in the standard mycobacterial cultivation media Middlebrook 7H9 or in modified LPB medium without additional sucrose (Fig. S7C). Importantly, investigating other mycobacterial strains under these conditions also revealed the presence of CWD cells, including the vaccine strains *Mycobacterium bovis* BCG Pasteur and BCG Russia, as well as *Mycobacterium marinum* M (Fig. 4C, 4D, S9). As shown in the phylogenetic tree (Fig. 4E), this indicates a widespread mycobacterial propensity to be able to form CWD cells in the presence of elevated concentrations of cell wall-targeting antibiotics.

**Figure 4.**
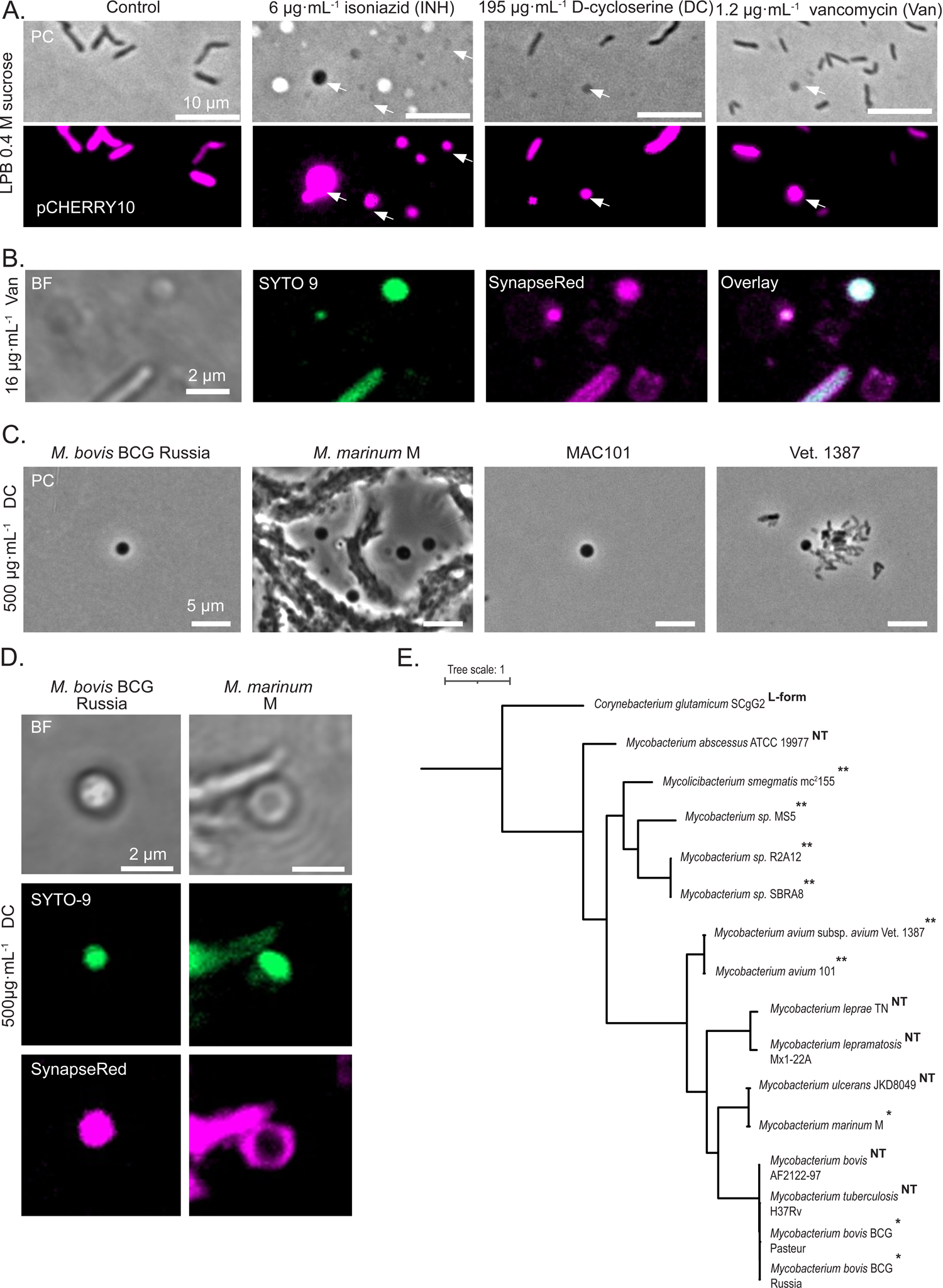
Cell wall-targeting antibiotics induces wall-deficiency in mycobacteria. **A)** Images of *M. smegmatis* mc^2^155::pCHERRY10 after 7 days of growth in the presence of 6 µgꞏmL^-1^ isoniazid, 195 µgꞏmL^-1^ D-cycloserine or 1.2 µgꞏmL^-1^ vancomycin. The control shows growth without the addition of antibiotics. CWD cells are indicated by arrows. Scale bars represent 10 µm. **B)** Confocal images showing CWD cells of *M. smegmatis* mc^2^155 formed after 5 days of growth in LPB (0.4 M sucrose) supplemented with 16 µgꞏmL^-1^ vancomycin. Nucleic acids and cell membranes were labeled with SYTO 9 and SynapseRed, respectively. CWD cells are indicated by arrows. Scale bars represent 5 µm. **C)** Phase contrast micrographs of *M. bovis* BCG Russia, *M. marinum* M and *M. avium* strains MAC101 and Vet. 1387 exposed to 500 µgꞏmL^-1^ D-cycloserine in LPB 0.4 M sucrose for 8 days. Scale bars represent 5 µm. **D)** Confocal images of DNA containing CWD cells of *M. bovis* BCG Russia and *M. marinum* M, formed after 8 days of growth in LPB (0.4 M sucrose) supplemented with 500 µgꞏmL^-1^ D-cycloserine. Nucleic acids and cell membranes were labeled with SYTO 9 and SynapseRed, respectively. Scale bars represent 2 µm. **E)** Phylogenetic tree of mycobacteria based on whole genome sequences. Asterisk(s) indicate strains tested in this study with the natural ability to form CWD cells in the absence (**) or in the presence (*) of antibiotics. Strains that were “not tested” are indicated with the designation NT. As outgroup *Corynebacterium* was included, which was shown to be capable of propagating as an L-form (Mercier *et al.,* 2014). Reference genomes are listed in Supplemental Table 6.

### Mycobacterial wall-deficient cells are not detected by conventional TB diagnostics

Our experiments show that several mycobacteria can form wall-deficient cells under hyperosmotic conditions and that their formation is stimulated by antibiotics. Although CWD cells have been reported in mycobacteria previously, the poor reproducibility of such studies has limited further investigations relating to their clinical relevance. Given that we were reproducibly able to generate and detect CWD cells in a range of mycobacteria, we set out to test whether current cultivation techniques and conventional diagnostics are appropriate to detect CWD cells. Noteworthy, Middlebrook medium (MB7H9), which is the standard cultivation medium for mycobacteria, does not sustain wall-deficient cells (Fig. 5A). However, in line with the previously used LPB medium, the addition of sucrose was sufficient to sustain such wall-deficient cells at a minimum sucrose concentration of 0.2 M (Fig. S10A). To determine the Minimal Osmolar Protection (MOP), the minimal osmolality needed to overcome turgor pressured exploding lysis through osmosis, the osmolality of these conditions was measured (Fig. S9B; Table S7). Interestingly, standard MB7H9 has an osmolality of 144 ± 1 mOsmꞏkg^-1^, which is below the minimum of 348 ± 1.2 mOsmꞏkg^-1^ that is needed to sustain wall-deficient cells (Fig. 5B).

**Figure 5.**
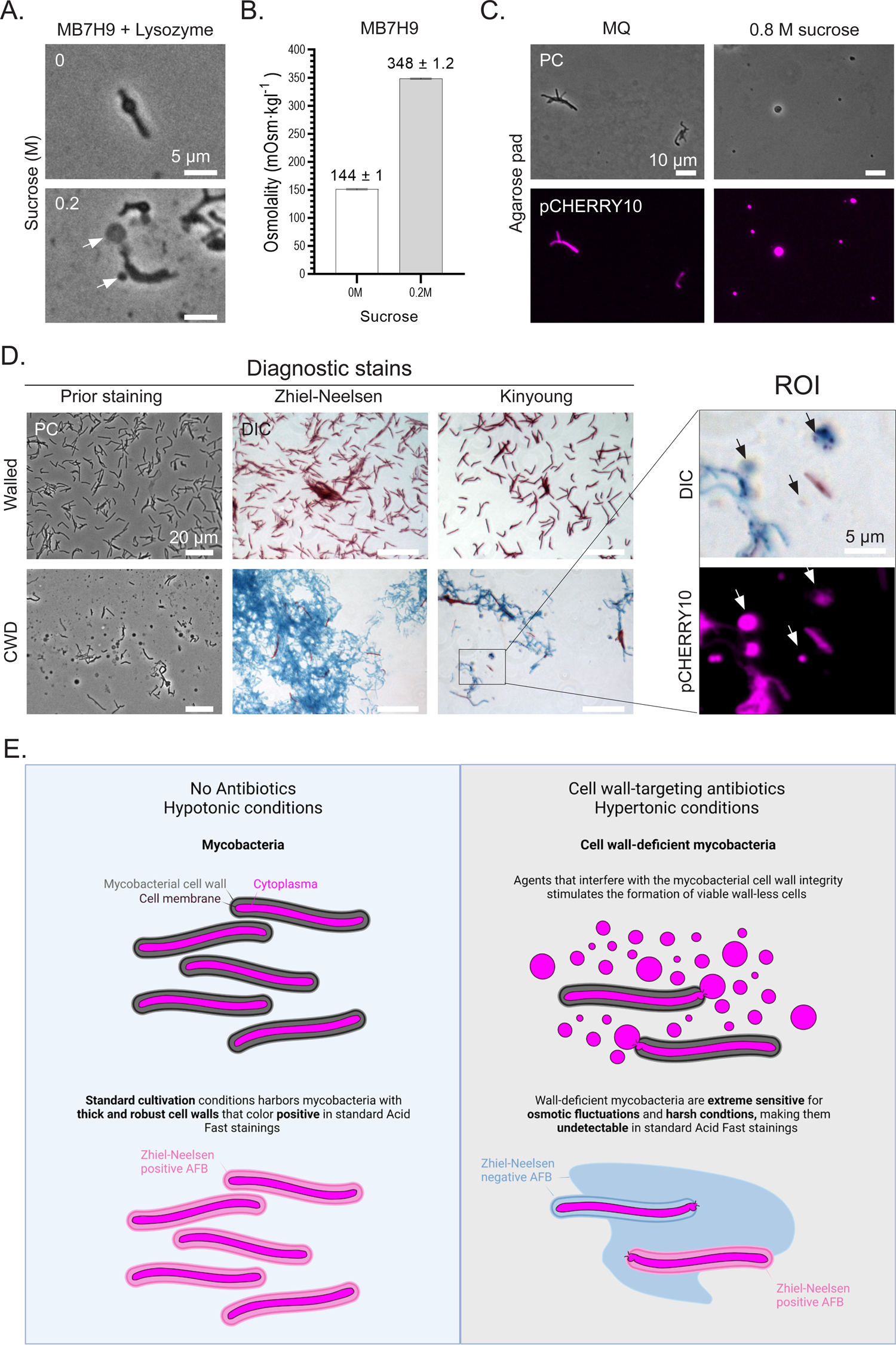
CWD cells of mycobacteria are undetectable by conventional diagnostics. **A)** Phase contrast images of *M. smegmatis* mc^2^155 after exposure to 1 mgꞏmL^-1^ lysozyme for 24 hours in MB7H9 supplemented with or without 0.2 M sucrose. The presence of CWD cells are indicated by arrows. Scale bars represent 5 µm. **B)** Osmolality measurements of MB7H9 supplemented with or without 0.2 M sucrose. Measurements were performed in sextuplicate. **C)** Images of *M. smegmatis* mc^2^155::pCHERRY10 CWD cells formed in LPB (1.0 M sucrose) mounted on MQ and MQ/sucrose (0.8 M) agar pads. Please note that all CWD cells are absent when placed on MQ agar pads. Scale bars represent 10 µm. **D)** DIC images of conventional Ziehl-Neelsen staining and Kinyoun staining on *M. smegmatis* mc^2^155::pCHERRY10 walled and enriched CWD cells. Acid fast bacteria (AFB) stain (positively) red, while the blue color indicates “background”. Phase contrast images were taken before the staining procedure. Please note that the fluorescent (CWD) cells are visible in the region of interest (ROI), which, however, do not stain red. CWD cells are indicated by arrows. Scale bars represent 10 and 5 µm. **E)** Schematic model representing the application of Ziehl-Neelsen staining on walled and CWD mycobacteria in hypo-(left) and hypertonic (right) conditions. Please note that wall-targeting agents further stimulate the formation of CWD cells, which remain undetected by this staining method. Created with <u>BioRender.com</u>.

These results emphasize the necessity of increasing the osmolality in standard mycobacterial cultivation media to detect cell wall-deficient cells, which should also be considered when visualising cells using common microscopy approaches (Fig. 5C, S9C). For instance, typically used fixation methods appear too harsh for CWD cells, as revealed in samples treated with conventional Acid-Fast Bacterial (AFB) staining methods (Fig. 5D). Notably, we show that the standard Ziehl-Neelsen hot staining method, which incorporates carbol fuchsin in the cytoplasm by heat application, led to destruction of all CWD cells. More specifically, samples containing mixtures of walled and CWD cells revealed abundant background staining but no CWD cells were found. To test whether the boiling steps in the staining procedure affect the preservation of CWD cells, we also applied the cold Kinyoun stain (Kinyoun 1915). This staining appeared to preserve the morphology of CWD cells, including their fluorescence (Fig. 5D ROI). However, these cells did not stain red as AFB-positive bacteria as would be expected from mycobacteria. In summary, our experiments reveal that mycobacterial CWD cells are not sustained on routinely used media and remain undetectable by conventional diagnostics (Fig. 5E).

## DISCUSSION

Conventional diagnostics continue to play a pivotal role in clinical settings in terms of detection and identification of pathogenic mycobacteria. In this study, we show that a wide range of mycobacteria undergo morphological transitions to form viable CWD cells in response to hyperosmotic stress and cell wall-targeting antibiotics. Contrary to mycobacterial extracellular vesicles (Prados-Rosales *et al*., 2011; 2014; Gupta & Rodrigues 2018), these CWD cells are larger in size, metabolically active and replication competent. Importantly, the conventional Middlebrook medium, which is used in mycobacterial growth indicator tubes (MGIT) and antibiotic susceptibility testing (Pfyffer *et al*., 1997), is not suitable to sustain mycobacterial CWD cells. Furthermore, traditional Acid-Fast Bacterial staining methods, such as Ziehl-Neelsen, also fail to detect mycobacterial CWD cells. This demonstrates that wall-deficient mycobacteria will not be detected using these classical approaches. Whole-genome sequencing (WGA) has recently been implemented in the diagnostics field, revolutionizing TB detection (Pankhurst *et al.,* 2016). While not experimentally tested here, the procedure of pre-processing the samples includes human cell lysis using osmotic shock (Votintseva *et al.,* 2017), which almost certainly would also destroy CWD cells. Taken together, given that conventional diagnostics are inappropriate to detect CWD cells, they may have been overlooked in clinical settings.

The observed shedding of the cell wall appears to be a general mycobacterial stress response, which is stimulated by the presence of antibiotics interfering with peptidoglycan and mycolic acid synthesis. The antibiotics used here included D-cycloserine and isoniazid, respectively, which are typically used in the treatment of (multi-drug resistant) *Mycobacterium tuberculosis* infections (Banerjee *et al.,* 1994; Hwang *et al.,* 2016). Since the closely related vaccine strain *M. bovis* BCG Russia (Borsch *et al*., 2007) was also shown to form CWD cells in response to these antibiotics, it raises the question if wall-deficiency plays a role in pathogenesis and persistence of *M. tuberculosis.* The formation of such cells may have further contributed to the emergence of multi-, extensively-, and totally drug-resistant (MDR/XDR/TDR) TB strains (Levin-Reisman *et al*., 2017), given that such wall-deficient cells are more competent for DNA uptake (Kapteijn *et al.,* 2022). The involvement of CWD cells in persistent infections could also explain the necessity of synergistical administration of cell wall-targeting antibiotics with agents that target other essential cellular processes, such as rifampicin (WHO, 2020).

Although the association of cell wall-deficiency in a wide range of infectious diseases has been suggested repeatedly, their potential role in chronic or in reoccurring infections remains controversial (Domínguez-Cuevas *et al*., 2012). More specifically, a CWD state of mycobacteria within the host has been proposed numerous times in clinical *in vivo* and *in vitro* studies (Markova, 2012; Mattman, 1970; Ratnam & Chandrasekhar 1976; Slavchev *et al*., 2016). However, these studies often lack accurate characterization and therefore remain largely inconclusive (Almenoff *et al*., 1996; Allan 2009). Therefore, cell wall-deficiency has been traditionally neglected within the mycobacterial field due to difficulties in reproducing and verifying results. In our work, we gained novel insights into the osmotic requirements of these mycobacterial CWD cells, providing reproducible protocols to generate and study this extreme form of morphological plasticity in mycobacteria. Taken together, it would be worthwhile to reinvestigate this phenomenon in terms of treatment persistence to determine its role in mycobacterial pathogenesis and antibiotic resistance.

Mycobacteria, like other actinobacterial relatives, incorporate newly synthesized cell wall material at the cellular tips, resulting in apical elongation, via a process known as polar growth (Howell and Brown, 2016). Disrupting the process of mycobacterial cell wall synthesis results in the extrusion of CWD cells at the polar tips. This is in line with recent observations in filamentous actinobacteria, in which hyperosmotic stress exposure results in apical extrusion of wall-deficient cells (Ramijan *et al.,* 2018). Since the formation of CWD cells is observed across the mycobacterial phylogenetic tree, it is conceivable that other mycobacteria are also capable of forming CWD cells such as *Mycobacterium abscesses*, *Mycobacterium ulcerans* and *M. tuberculosis* (Fig. 4E). Remarkably, *M. marinum* and *M. bovis* BCG strains only formed CWD cells in response to cell wall-targeting antibiotics, indicating that the integrity of the cell wall of these strains is intrinsically stronger compared to other non-tuberculosis mycobacteria (NTM). Perhaps this increased cell wall integrity is due, at least in part, to the presence of methyl-branched fatty acids-containing lipids, such as phthiocerol dimycocerates (PDMIs), that are unique to these pathogenic mycobacteria (Yu *et al.,* 2012). The role of increased cell wall integrity, however, in terms of mycobacterial morphological plasticity and virulence needs to be further elucidated.

In conclusion, our work shows that mycobacteria form viable CWD cells in response to environmental stressors and that this response is reversible. CWD cells are undetectable by classical diagnostic approaches and in all likelihood have therefore been over-looked in clinical settings. It is advisable to revisit current diagnostic approaches to ensure proper detection of CWD cells. Potentials in future detection of mycobacterial CWD cells could, for example, involve targeting mycobacterial specific plasma membrane components, such as phosphatidylinositol dimannosides (PIM2) (Sohlenkamp and Geiger, 2016). As some of the most important antigens of mycobacteria are cell wall-associated glycolipids and lipoproteins, the loss of the cell wall would potentially allow such cells to escape from recognition by the host immune system, further contributing to host persistence (Källenius *et al*., 2016). Their role in mycobacterial pathogenesis deserves more in depth attention. If indeed CWD cells are a major contributor to bacterial persistence, further studies are needed to investigate their role in the pathogenesis of (chronic and recurring) mycobacterial infections, as well as to provide important new leads for discovery and development of better antimicrobial agents to combat the devastating diseases caused by these bacteria.

## MATERIAL AND METHODS

### Mycobacterial strains and culture conditions

All Mycobacterium used in this study are listed in Table S1. Unless stated otherwise, the strains were cultured on Middlebrook 7H10 agar (BD Difco) supplemented with 0.5% (vol/vol) glycerol (Duchefa G1345) and 10 % (vol/vol) Middlebrook oleic acid-albumin-dextrose-catalase (OADC; BD BBL) added before and after autoclavation, respectively. For growth in liquid, the strains were cultivated in Middlebrook 7H9 broth (BD Difco) supplemented with 0.2% (vol/vol) glycerol and 10 % (vol/vol) albumin-dextrose-catalase enrichment (ADC; this lab) added before and after autoclavation, respectively. The ADC enrichment was made following the composition of the BD BBL ADC enrichment, containing per litre 8.5 g sodium chloride (Sigma-Aldrich 31434), 50 g Bovine Serum Albumin heat shock fraction (BSA; Sigma-Aldrich A9647), 20 g D-glucose (Duchefa G0802) and 0.03 g Catalase from bovine liver (Sigma-Aldrich C9322). The plasmid pCHERRY10 was obtained from Addgene (#24664), to fluorescently label *M. smegmatis* mc^2^155 with mCherry under the constitutively G13 promoter. To culture the fluorescent strain, the antibiotic hygromycin B (Duchefa) was added to a final concentration of 100 µgꞏmL^-1^. *M. marinum* and mycobacterial endophytes were cultivated at 30 °C at 200 rpm and 100 rpm, respectively. The remaining mycobacterial strains were grown at 37 °C, with an agitate on speed of 200 rpm. To obtain exponential growing cells of *M. smegmatis* mc^2^155, the strain was inoculated from a colony and precultured in 10 mL MB7H9 (Erlenmeyer flask; 50 mL) to stationary phase. This preculture was diluted 1000-fold in 50 mL MB7H9 in 250 mL coiled Erlenmeyer flasks, resulting into exponential grown cells after 16 - 24 hours, with an OD_600nm_ = 0.4 – 1.0.

To support growth of mycobacterial CWD cells, the strains were grown in media previously used for cultivating CWD cells of filamentous actinobacteria (Ramijan *et al*. 2018). Adjusted solid L-Phase Medium Agar (LPMA) contains 0.5 % D-glucose (w/v), 0.5 % Yeast Extract (w/v) (Gibco™), 0.5 % Bacto^™^ Peptone (w/v) (Gibco™), 0.58 M Sucrose (w/v) (Duchefa S0809), 0.01 % MgSO4ꞏ7H2O (w/v) (Sigma-Aldrich 10034-99-8) and 0.75 % Iberian agar (w/v), supplemented with 5 % Horse Serum (v/v). Standard adjusted liquid L-Phase Broth (LPB) medium is composed of yeast extract (w/v), 0.25 % Bacto^™^ Peptone (w/v), 0.15 % Malt Extract (w/v) (Duchefa M1327), 0.5% D-glucose (w/v), 1.5% Bacto^™^ Tryptic Soy Broth (w/v) (BD^™^ 211825) and 0.64 M Sucrose. Cultures were incubated either at 30 °C or 37 °C, while slowly shaking at 100 rpm.

To investigate the effect of the sucrose concentration on bacterial growth and extrusion of CWD cells, LPB medium with different amounts of sucrose were prepared (ranging from 0-1.0 M sucrose). For the detection of CWD cells, *M. smegmatis* Mc^2^155::pCHERRY10 was grown to exponential phase, as described, and inoculated in Erlenmeyer flasks containing 10 mL LPB supplemented with the sucrose gradient to an high optical density (final OD_600nm_ = 0.1) in duplicate. After 16 hours, the presence of CWD cells was analysed by phase contrast microscopy and quantified by flow cytometry, as described below. To assess bacterial growth, the same exponential phase grown Mc^2^155::pCHERRY10 was inoculated with low turbidity (final OD_600nm_ = 0.01) in a flat polystyrene 96-well plate ([GRE96ft] - Greiner 96 Flat Transparent Cat. No.: 655101/655161/655192) containing 200 µL standard LPB supplemented with sucrose in triplicates. Growth output was assessed through OD_600nm_ measurements in technical duplicates using a TECAN Spark^®^ 10M with SparkControl (v 2.3) software.

### Enrichment of CWD cells

To enrich for mycobacterial CWD cells, *M. smegmatis* was grown to exponential phase as described, inoculated to an high cell density (OD_600nm_ = 0.1) in 100 mL standard LPB or LPB with 1.0 M sucrose and incubated at 37 °C, while shaking at 100 RPM. CWD cells were filtered through a 11 µm pore size cellulose Whatman™ filter (Sigma Aldrich, WHA1001055), utilizing a 47 mm Magnetic Filter Funnel (VWR 516-7597) and a vacuum pump, and enriched by centrifugation of the supernatant for 1 hour at 1000 x g. Finally, cells were resuspended into 1-5 mL of the remaining supernatant.

### Generation of protoplasts

To obtain mycobacterial protoplasts, a protocol was developed based on existing protocols used to generate protoplasts in *Streptomyces* (Kieser, 2000) and spheroplasts in Mycobacteria (Udou, 1982). In short, cells were grown to exponential phase (OD_600nm_ = 0.4) in standard MB7H9, after which 1 % glycine (Duchefa) was added followed by growth for another 16-24 hours. Cells were harvested at 4000 x g for 10 min and 2x washed at 1200 x g for 14 min in Sucrose Magnesium (SM) solution (10.3 % sucrose (w/v) and 25 mM MgCl_2_ (Duchefa M0533) in MQ). Cells were then resuspended in Lysozyme solution (pellet of 20 mL starting culture in 4 mL solution), containing 1 mgꞏmL^-1^ Lysozyme (Sigma-Aldrich 62971) in protoplast buffer as described in Kieser, and incubated at 37 degrees while standing for 16-20 hours. Briefly, basic protoplast buffer is prepared by adding 10.3 g Sucrose, 25 mg K_2_SO_4_, 0.202 g MgCl_2_ꞏ6H_2_O and 200 µL trace elements in 80 mL MQ and autoclave for 20 min at 120 °C. Trace elements consist of (L^-1^) 40 mg ZnCl_2_, 200 mg FeCl_3_ꞏ6H_2_O, 10 mg CuCl_2_ꞏ2H_2_O, 10 mg MnCl_2_ꞏ4H_2_O, 10 mg Na_2_B_4_O_7_ꞏ10H_2_O and 10 mg (NH_4_)_6_Mo_7_O_24_ꞏ4H_2_O. Protoplast buffer then is enriched with 1 mL KH_2_PO_4_ (0.5%), 10 mL CaCl_2_ꞏ2H_2_O (5M, 3.68%) and 10 mL TES buffer (5.73%, 0.25 M, pH 7.2). Successful protoplasting was monitored using phase contrast microscopy. After protoplasts were formed, cell debris was removed by filtration through a SM prewashed 40 µm cell strainer (PluriStrainer® SKU 43-50040-51).

### Light microscopy

To immobilize cells, unless stated otherwise, basic LPM agar pads lacking horse serum were made utilizing a 15 mL falcon tube on a thin layer agar plate. Alternative, ultrapure MilliQ (MQ) water containing 1.5 % agarose with or without supplementation of 30 % sucrose was used to generate pads. For imaging, 5 µL of sample was either loaded on an agar pad placed on a glass slide (Fisher 1157-2203) and covered with 20 x 20 mm (Epredia #1) coverslip, or directly loaded in a 35 mm imaging µ-dish (Ibidi®) and covered by an agar pad. Phase contrast images were obtained using a Zeiss Axio Lab A1 upright microscope equipped with a Zeiss Axiocam 105 color (resolution 5 mega pixel, 2.2 µm/pixel) and collected using Zen 2 software (blue edition, Carl Zeiss Microscopy GmbH). Differential Interference Contrast (DIC), phase contrast and fluorescent images were taken with a Zeiss Axioplan 2 upright microscope equipped with an Axiocam Mrc 5 camera utilizing AxioVison Rel. 4.8.1.0 Zen software (Carl Zeiss Imaging Solutions GmbH). Fluorescent filter sets applied were 63 HE (Carl Zeiss, consisting of a 572/25 bandpass excitation filter, 590nm beam splitter, and 629/62 nm bandpass emission filter) to capture mCherry fluorescence. Fluorescent live imaging was performed on a Zeiss AXIO Observer.Z1 equipped with a Hamamatsu ImagEM X2 EM-CCD camera C9100 utilizing Zen 2.6 software (blue edition, Carl Zeiss Microscopy GmbH). Confocal images were acquired with a Zeiss LSM 900 confocal microscope with Airyscan 2 module, temperature control chamber and Zen 3.1 software (blue edition, Carl Zeiss Microscopy GmbH). All excitation and emission settings for this microscope are listed in Supplemental Table S2. Microscopy images were processed using either OMERO or ImageJ/FIJI (Schindelin *et al*., 2012).

### Fluorescent probes

Fluorescent probes were added to 25 µL aliquots of liquid cultures or resuspended colonies, followed by incubation at room temperature for 15 min. Nucleic acids were stained using 2 µM SYTO 9 (S34854, Invitrogen). The plasma membrane was labelled using SynapseRed C2M (SynapseRed, PK-CA707-70028, PromoKine, PromoCell GmbH; also known as FM5-95, a trademark of Molecular Probes, Inc.) to a final concentration of 70 µM.

### Cell diameter measurements

Cell diameters of extruded CWD *M. smegmatis* mc^2^155 cells, grown in LPB supplemented with 1.0 M sucrose in triplicate, were measured overtime by fluorescent labelling of cell membranes with SynapseRed through confocal micrograph acquisitions. Spherical structures that were not positively labelled with SYTO 9, and therefore lacking nucleic acids, were excluded from the assessment. Micrographs were analysed using FIJI/ImageJ (Schindelin *et al.,* 2012).

### Flow cytometry

To quantify walled and CWD cells, flow cytometry was used. Briefly, red fluorescent *M. smegmatis* mc^2^155::pCHERRY10 cells were filtered through a LPB prewashed 40 µm cell strainer (PluriStrainer® SKU 43-50040-51) prior to quantification using a Bio-Rad S3e Fluorescence-activated Cell Sorter (FACS) equipped with ProSort™ Software, version 1.6. Fluorescence was detected following excitation at 561 nm and by using a 615/25 nm bandpass emission filter (FL3). As sheath fluid, ProFlow Sort Grade 8x Sheat Fluid (Bio-Rad Laboratories, inc; #12012932) was used. Quality control (QC) was passed utilizing ProLine™ Universal Calibration Beads (Bio-Rad Laboratories, inc; #1451086). The threshold was set on 0.02 FCS to minimize background noise and the flow was adjusted to 500 events per second. For the sucrose gradient 50,000 events were collected. For the triton viability assay, 20,000 events were collected and experiments were performed in triplicate. All data analysis was done using FlowJov10.8.1 (BD BioSciences). To remove outliers, three successive gates were drawn in 1.1ꞏ10^0^ Side Scatter (SSC) x 1.1ꞏ10^0^ Forward Scatter (FSC) Height, 1.5K - 2.5K SSC Width x Height and 1.5K-2.5K FSC Width x Height, respectively (Fig. S2). iQuadrant bins were drawn to roughly distinguish walled and CWD cells, typically 2ꞏ10^1^ SSC Height over 1ꞏ10^1^ FL3 (Sucrose gradient & Protoplasts) and 5ꞏ10^1^ SSC Height over 4ꞏ10^1^ FL3 (Triton viability) to select for complexity and fluorescence. Specific gates were drawn to include populations of interest in SSC x FL3, including subpopulations in lower sucrose conditions of propagating walled cells. Relative population percentages represented by the populations “Walled fluorescent” and “CWD fluorescent” were not further utilized for analysis. Biological and technical replicates were separately processed for graphical representation and concatenated for statistical analysis and visualization purposes. Pearson Chi-squared were performed on detergent treatment compared to expected no treatment within separate iQuadrant bin populations (df = 2) or over the total iQuadrant bins population (df = 4), as found in Supplemental Table 3.

### Cryo-Transmission Electron Microscopy

Mycobacterial cells were prepared and checked by phase contrast for cell morphology and cell density. From the prepared cells, 20 µL samples were mixed with 2 µL 10 nm colloidal gold beads (Protein A coated, CMC Utrecht), of which 3-4 µL was applied on glow-discharged 200 mesh copper grids with an extra thick R2/2 carbon film (Quantifoil Micro Tools). Vitrification was performed using an automated Leica EM GP plunge-freezer, which automatically pre-blots for 30 seconds, followed by 1 second blotting step and plunge-freezing of the grid into liquid ethane. The samples were mounted on a 626 cryo-specimen holder (Gatan, Pleasanton, CA) and imaged using a 120 kV Talos TEM (Thermo Fisher Scientific) equipped with a Lab6 electron emission source and Cita detector (Thermo Fisher Scientific). The resulting Cryo-TEM micrographs have a pixel size of 0.34 nmꞏpixel^-1^. The density plots of the mycobacterial cell envelope were generated through FIJI/ImageJ (Schindelin *et al.,* 2012).

### Viability assay

To test the viability of mycobacterial CWD cells, the pCHERRY10 expressing *M. smegmatis* mc^2^155 cells were exposed to the detergent Triton x-100 (TX-100) (PanReac AppliChem) and plated on MB7H10 medium to detect colony forming units (CFU). CWD cells were generated as described and enriched after three days of growth. As a walled control, the cells were grown in MB7H9 to exponential phase in 24 hours. As a control, protoplasts were generated as described above. The cells were then exposed for 15 min to either MilliQ (MQ) or 0.1% MQ + TX-100. To detect viable CFUs, the samples were 10-fold diluted in LPB (20 µl in 180 µl) in polystyrene 96-well plate utilizing multichannel pipets. 10 µl spots of diluted samples were plated in duplicate on MB7H10, dried for 15 min and incubated for 5 days at 37 °C. After spotting, the cells were inspected through flow cytometry and DIC-fluorescence for the presence of spherical mCherry-expressing cells. Paired student t-tests were performed were performed on TX-100 treatment compared to no treatment, as found in Supplemental Table 4.

### Antibiotic CWD induction assay

Minimal inhibitory concentration (MIC) assays were performed to detect their effect on the formation of CWD cells. For this, 2-fold dilutions of antimicrobial agents in MB7H9, LPB 0 M sucrose and LPB 0.4 M sucrose were prepared in polystyrene 96-wells plate ([GRE96ft] - Greiner 96 Flat Transparent Cat. No.: 655101/655161/655192), including starting concentrations (w/v): 50 mgꞏmL^-1^ isoniazid (Duchefa), 50 mgꞏmL^-1^ D-cycloserine (Duchefa) and 5 mgꞏmL^-1^ vancomycin (Duchefa). MIC assays were performed in triplicate and contained a Blanco measurement per concentration. Exponential growing *M. smegmatis* mc^2^155::pCHERRY10 cells were inoculated at a turbidity of OD_600nm_= 0.01, incubated 100 RPM for 7 days. Incubation was performed at 30 °C and wrapped in parafilm to reduce well evaporation. Output was assessed through turbidity measurements using a TECAN Spark^®^ 10M with SparkControl (v 2.3) software and by scanning the plates. Based on the MICs, antibiotic CWD induction assays were reproduced in Erlenmeyer flasks containing 10 mL LPB with 0 or 0.4 M sucrose supplemented with antibiotics, including 16 µgꞏmL^-1^ isoniazid, 16 µg/mL vancomycin and 400 µgꞏmL^-1^ D-cycloserine. Controls included supplementation with either 1 mgꞏmL^-1^ Lysozyme + 25 mM MgCl_2_ (positive control) or no supplementation (negative control). To test other mycobacterial species, all the antibiotic concentrations were increased to 500 µgꞏmL^-1^ to overcome species-specific antibiotic MICs and to ensure antibiotic effectiveness. All CWD induction assays were inoculated in high turbidity of OD_600nm_= 0.1 and grown 5-8 days at 30 or 37 °C 100 RPM mL. Non-fluorescent strains were checked by phase contrast microscopy and subjected to fluorescent dyes to assess the presence of nucleic acids and membranes by confocal microscopy.

### Phylogenetic analysis

For DNA isolation of strains that had not been sequenced, colonies were harvested from fresh agar plates. Samples were washed 3 times with sterile PBS and snap frozen. DNA was subsequently isolated using the method of Bouso *et al*. (2019) without shearing. For the library/run mode, the PromethION SQK-LSK109 ligation sequencing kit was used. Samples were run on an ONT GridION by FG-technologies, Leiden, The Netherlands. Base calling was done using Gupy v4.0.11. The sequencing data are available under NCBI submission number PRJNA895223. Genome sequences were used to construct a maximum-composite likelihood phylogeny tree using PhyloPhlAn (Segata *et al.,* 2013). Ortholog identification and alignment was performed in Phylophlan using the “-u” command. The phylogenetic tree was visualized using iTOL (Letunic and Bork, 2007). Reference genomes are listed in Supplemental Table 6.

### Quantification of osmolality and osmotic pressure

To test conventional mycobacterial culture medium in their ability to sustain CWD cells, ascending sucrose amounts were added to conventional Middlebrook 7H9 and LPB media. Osmolality of the media was measured in sextuplicate using freezing point depression with an osmometer (Micro-Osmometer Autocal Type 13, Roebling), calibrated using distilled water (0) and 300 mOsm standard ampulla delivered with the instrument. To calculate the osmotic pressure (π) the following equation was applied: *π (mmHg) = 19.3 x osmolality (mOsm*ꞏ*kg^-1^)* (Rasouli, 2016). The mean values are listed in Table S7. To test the sustainability of CWD cells, *M. smegmatis* mc^2^155 was grown to exponential phase, inoculated in the media with high turbidity (OD_600nm_= 0.1) and exposed to 1 mgꞏmL^-1^ lysozyme and 25 mM MgCl_2_ for 24 hours at 37 °C at 100 rpm. The conditions were screened for the presence of spherical cells by phase contrast microscopy.

### Mycobacterial diagnostics

Conventional carbol fuchsin staining was performed using hot Ziehl-Neelsen complete staining kit (CLIN-TECH limited product code 62103, BioTrend) (Ziehl, 1882; Neelsen, 1883). In short, the stain was prepared as follows: 1) 100 µL of cells were spread and airdried on glass slides. 2) The cells were fixated by passing the glass slides 3 times through the flame. 3) The glass slides were flooded with carbol fuchsin (ZN) and heated above the flame until steam raised. 4) Slides were washed with demineralized water. 5) 3 % alcohol acid (hydrochloric acid) was applied until no changes in color occurred anymore. 5) The glass slides were rinsed with demineralized water and counter stained with 0.05 % methylene blue for 10 seconds. 6) Slides were rinsed with water, drained and airdried. 7) Slides were examined using oil immersion. Alternative carbol fuchsin staining was performed using cold Kinyoun staining (CLIN-TECH limited product code 621045, BioTrend) (Kinyoun, 1915). In short, this stain was prepared as follows: 1) 100 µL of cells were spread and airdried on glass slides. 2) The cells were fixated by passing the glass slides 3 times through the flame. 3) The glass slides were flooded with carbol fuchsin (Kinyoun’s) for 1 minute. 4) Slides were gently washed with demineralized water. 5) 3 % alcohol acid was applied until no changes in color occurred anymore, in less than 30 seconds. 5) The glass slides were directly rinsed with demineralized water and counter stained with 0.05 % methylene blue for 30 seconds. 6) Slides were rinsed with water, drained and air-dried. 7) Slides were examined with DIC microscopy using oil immersion.

## ACKNOWLEDGEMENTS

This work was supported by an NWO Open Competition ENW-Klein grant (OCENW.KLEIN.039) to D.C. We thank Wilbert Bitter for sharing *Mycolicibacterium smegmatis* mc^2^155, Annemarie Meijer for sharing *Mycobacterium marinum* M, Jos Raaijmakers for sharing the endophyte isolates, Alexandra Aubry (Sorbonne University, France) for sharing *Mycobacterium avium* human clinical isolate 1904038568, and Giovanni Ghielmetti (University of Zurch, Switzerland) for sharing the veterinary strains of *M. avium*.

## AUTHOR CONTRIBUTIONS

N.D. and T.W. carried out the experiments. V.C.B. constructed the phylogenetic tree, while H.P.S. sequenced the Mycobacterial genomes. All authors contributed to the experimental design and discussion of results. N.D and D.C wrote the manuscript with input from all authors.

## SUPPLEMENTAL FIGURES LEGENDS

**Figure S1.**
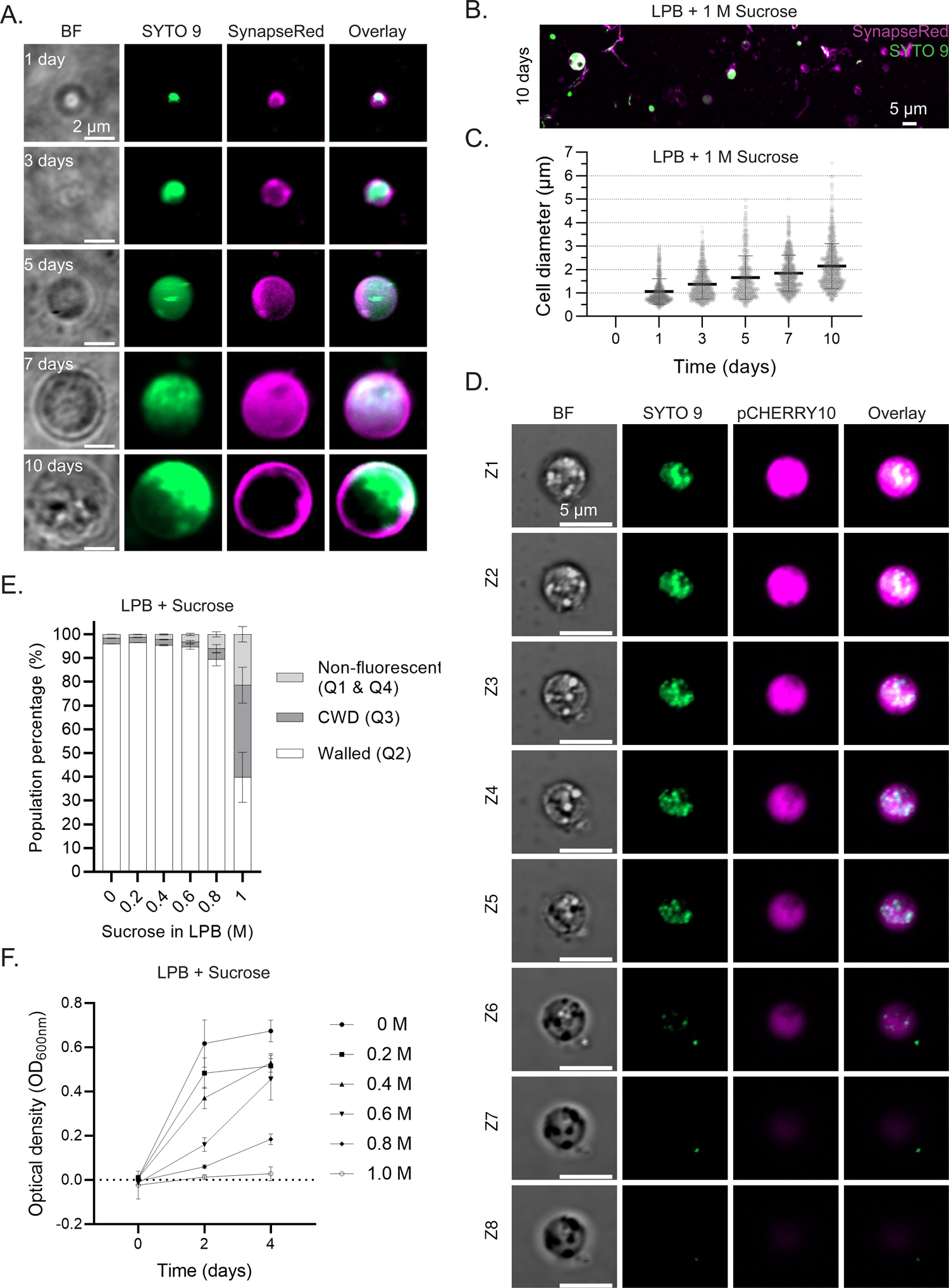
Formation of CWD cells in *M. smegmatis* containing DNA. A) Confocal images of nucleic acid containing CWD cells from *M. smegmatis* mc^2^155 with relative large diameters formed overtime in LPB. Nucleic acid and membrane were labeled with SYTO 9 and SynapseRed, respectively. Scale bars represent 2µm. B) Image of *M. smegmatis* mc^2^155 walled cells and CWD cells grown for 10 days in LPB (1.0 M sucrose). Nucleic acid and membrane were labeled with SYTO 9 and SynapseRed, respectively. Scale bar represent 5 µm. C) Cell diameter measurements of nucleic acid containing CWD cells over time (n = 480-684 per time point, single measurement per CWD cell). Graph represents measured means and standard deviation. D) Confocal Z-stack images of *M. smegmatis* mc^2^155::pCHERRY10 CWD cell grown 10 days in LPB. Nucleic acids were labeled with SYTO 9 for 15 min. Confocal images were taken in 8 times Z-stacks of 270 nm. Scale bar represents 5 µm. E) Quantification of flow cytometry data (see Fig. S2) of *M. smegmatis* mc^2^155::pCHERRY10 grown for 2 days in LPB containing various concentrations of sucrose. White bars represent walled cells, while light grey and dark grey represent non-fluorescent particles and CWD cells, respectively. Please note that increasing the concentration of sucrose results in a higher number of CWD cells. F) Growth curve based on optical density measurements (OD_600nm_) of *M. smegmatis* mc^2^155::pCHERRY10 grown for 4 days in LPB containing various concentrations of sucrose. Please note that increasing the concentration of sucrose, results in the inhibition of growth.

**Figure S2.**
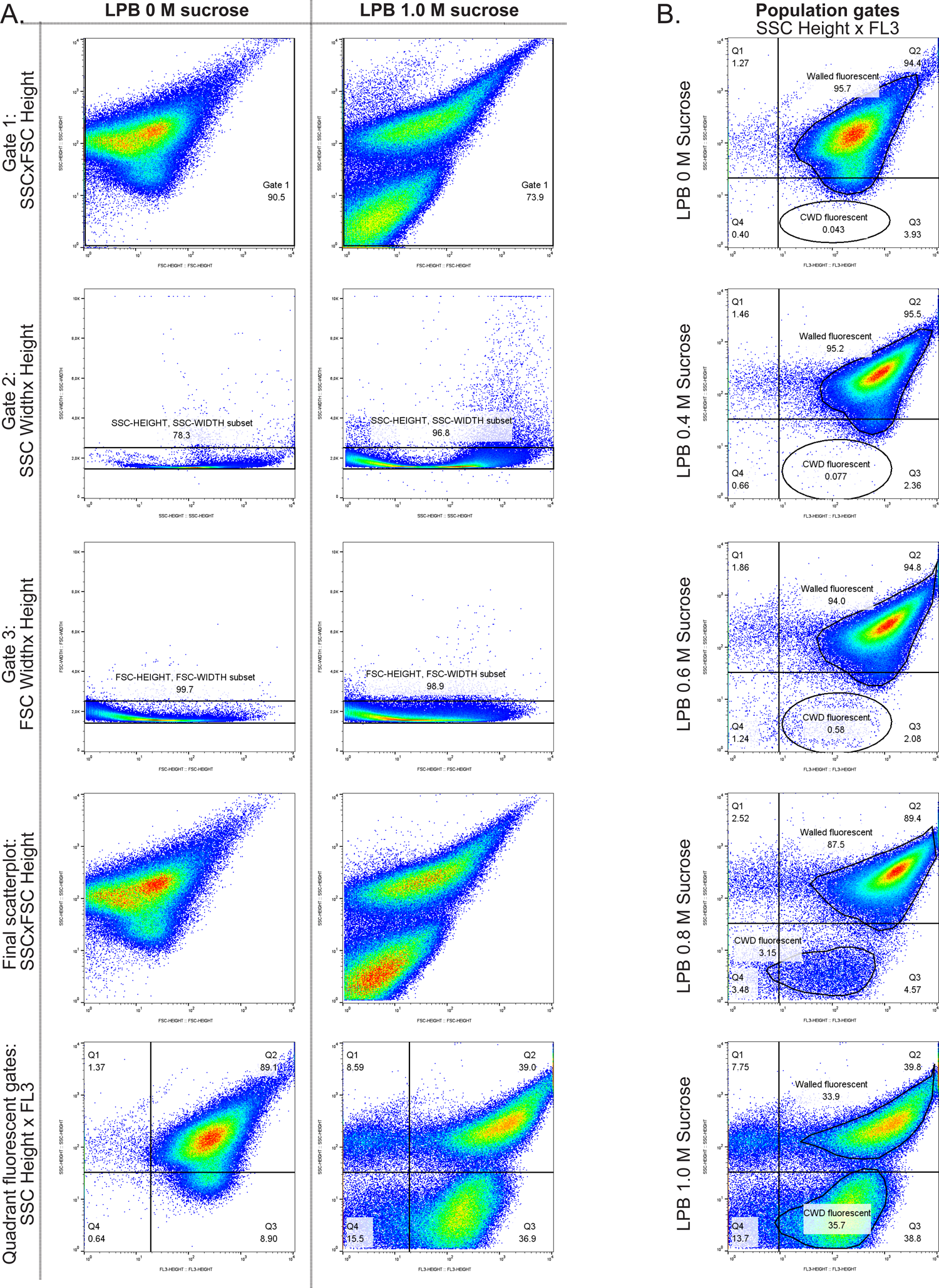
Flow cytometry processing of fluorescent labelled walled and CWD cells of *M. smegmatis* Flow cytometry gating workflow for remove outliers and select populations of interest. **A)** Removal of outliers represented by the two extremity conditions, LPB supplemented with 0 M and 1.0 M sucrose. Gate 1 in the FSC x SSC Height log, to remove background outliers present on both axis. Gate 2 in the SSC Height x Width log to remove outliers in the SSC. Gate 3 in the FSC Height x Width log to remove outliers in the FSC. Populations represented in SSC x fluorescent filter 3 (FL3), bandwidth 615/25 with laser 56.1 Simple iQuadrants bins gating for estimated separation of populations, namely Highly complex, non-fluorescent particles (Q1), Highly complex, fluorescent particles (Q2, Walled), Less complex, fluorescent particles(Q3, CWD cells), and Less complex, non-fluorescent particles (Q4). **B)** Specific gates drawn to specific populations of interest in SSC x FL3, including subpopulations in lower sucrose conditions of propagating walled cells with nascent, less complex, cell walls prior cell wall modifications. Relative population percentages represented by the populations “Walled fluorescent” and “CWD fluorescent”.

**Figure S3.**
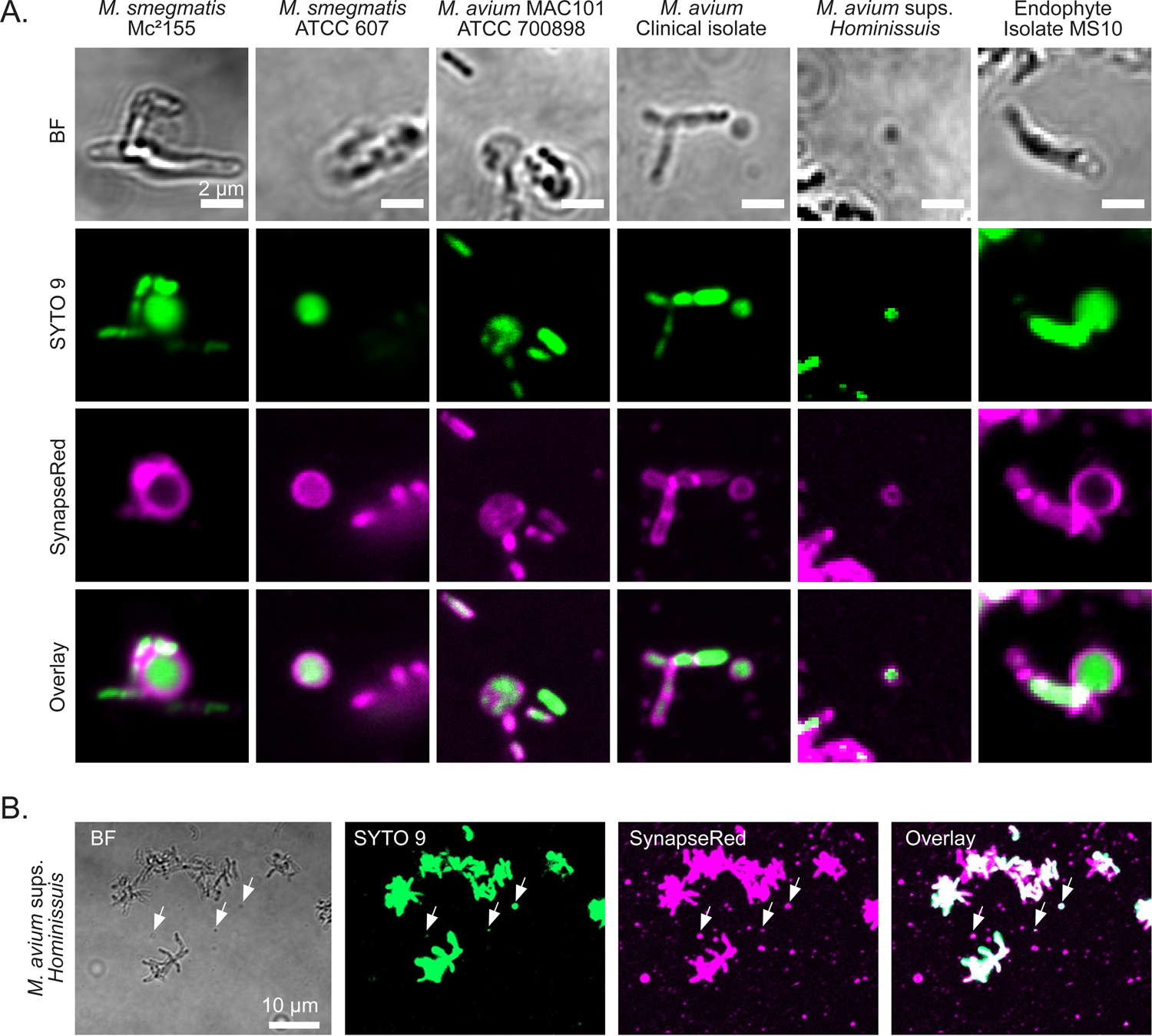
CWD formation in response to hyperosmotic conditions in various mycobacterial species Confocal micrographs of various mycobacterial strains grown in LPB (1.0 M sucrose). Cells were fluorescently dyed with SYTO 9 and SynapseRed to label nucleic acids and membranes, respectively. **A)** Mycobacterial strains were grown for either 2 days (*M. smegmatis* strains) or 2 weeks (*Mycobacterium avium* complex strains) at 37 °C 100 RPM, or were grown 1 week at 30 °C 100 RPM (endophytic isolates). Scale bars represent 2 µm. **B)** Veterinary isolate *Mycobacterium avium* subsp. *hominissuis* 20-935 /2 grown for 2 weeks at 37 °C 100 RPM. Please note that this strain produces some CWD cells, but mostly empty membrane vesicles. Scale bar represent 10 µm.

**Figure S4.**
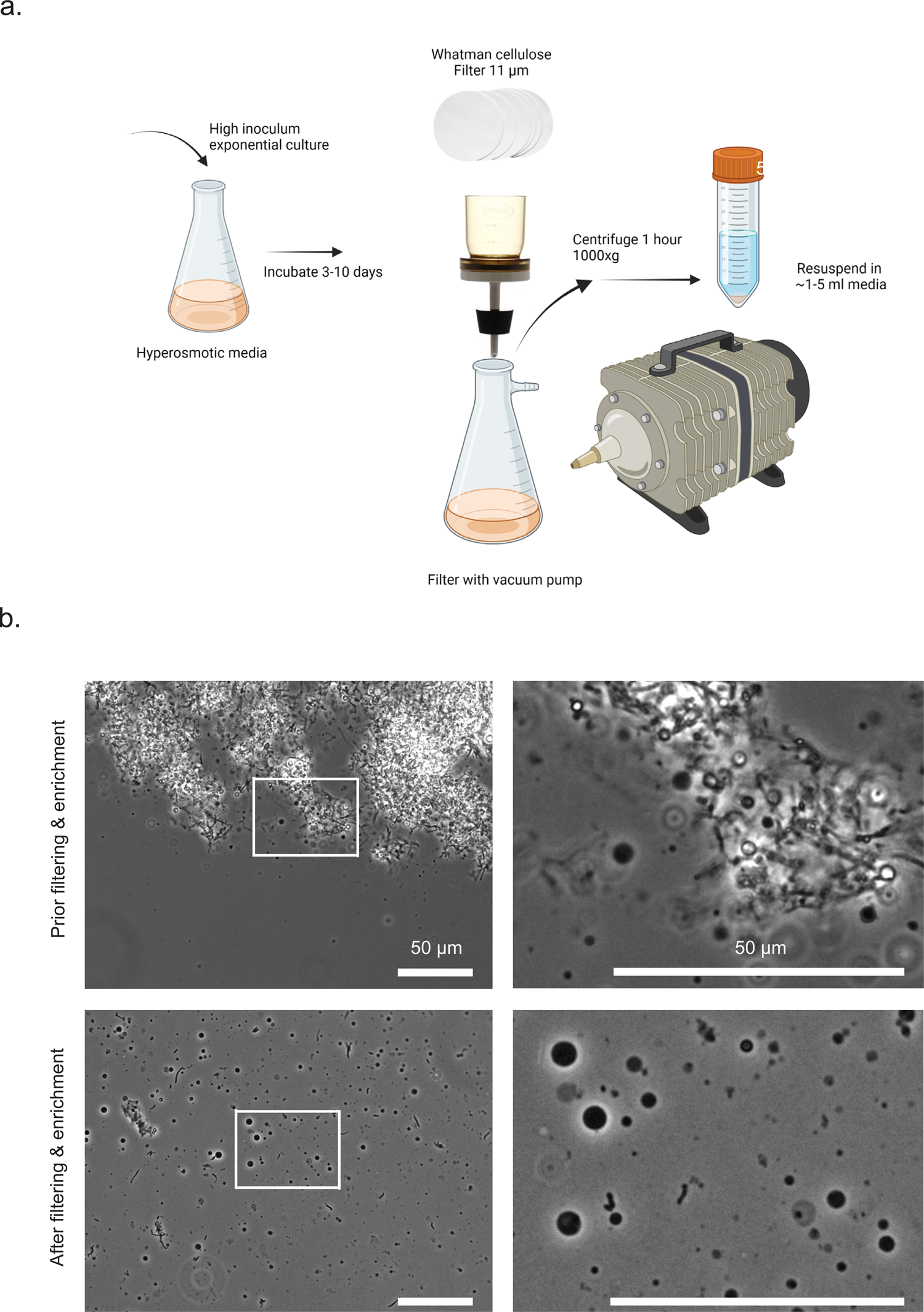
Enrichment of mycobacterial CWD cells. **A)** Enrichment method of CWD cells utilizing particle retention of 11 µm and slow centrifugation. In short, CWD cells are produced in as described in Material and Methods. Incubate until CWD cells are formed and filter culture with vacuum pump. Centrifuge filtrate for 1 hour at low centrifuging force (1000 x g). Please note that low centrifuging force is needed to avoid cell shearing of the sensitive CWD cells. Resuspend loose pellet in left-over supernatant. Created with <u>BioRender.com</u>. **B)** Phase contrast images of *M. smegmatis* mc^2^155 CWD cells grown for 8 days in LPB (1.0 M sucrose), before and after enrichment. Scale bars represent 50 µm.

**Figure S5.**
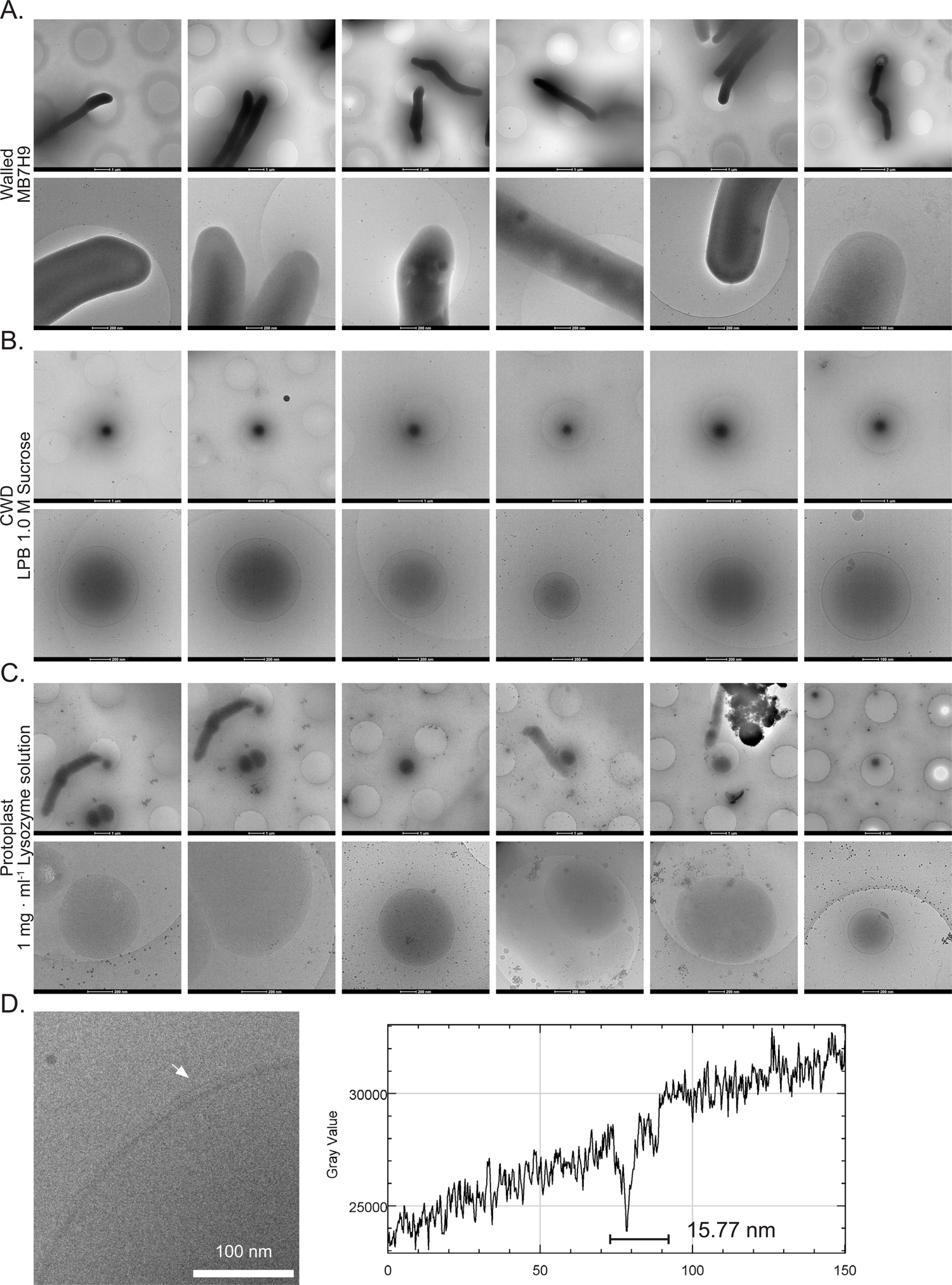
Cryo-TEM images of various cells of *M. smegmatis*. **A)** Cryo-TEM images of exponential-phase grown *M. smegmatis* mc^2^155 in MB7H9 liquid media, blotted directly from culture. Selected walled cells used for analysis. Scale bars represent 1 µm, 2 µm, 200 nm or 100 nm. **B)** Cryo-TEM micrographs of *M. smegmatis* mc^2^155 enriched CWD cells, grown 8 days in LPB 1.0 M Sucrose. Selected CWD cells used for analysis. Scale bars represents 1 µm, 200 nm or 100 nm. **C)** Cryo-TEM micrographs of *M. smegmatis* mc^2^155 artificial produced protoplasts in P-buffer. Selected protoplasts used for analysis. Scale bars represents 1 µm or 200 nm. **D)** Cryo-TEM micrograph region of interest in shows wall-like structures detected on top of plasma membrane of *M. smegmatis* mc^2^155 enriched CWD cell, indicated by arrow. Density plot of 150 nm across cell membrane shows an extra depth representing the wall-like structure, measuring an increase in cell-envelope size to 15.77 nm. Scale bar represents 100 nm.

**Figure S6.**
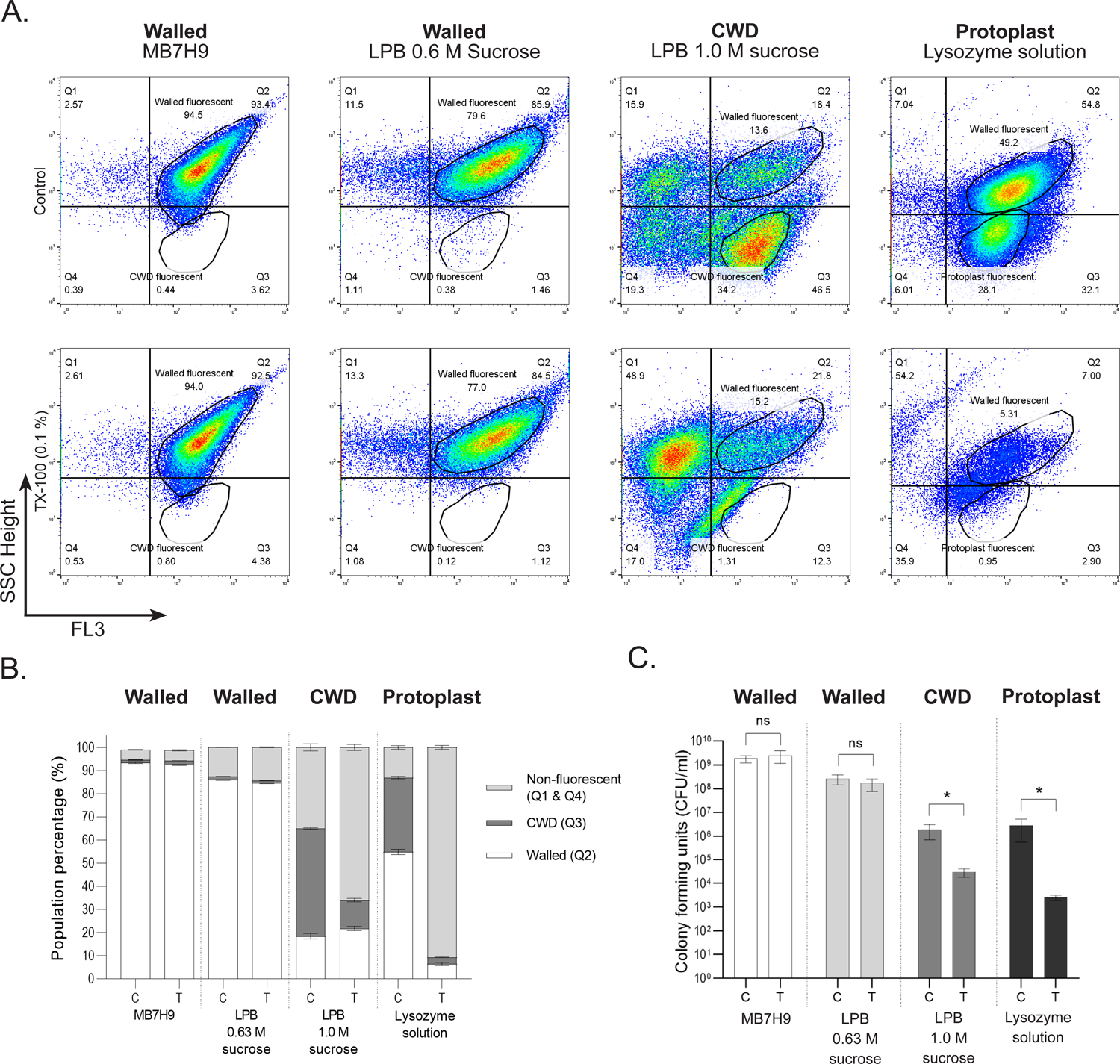
CWD cells are sensitive to detergent Triton X-100. **A)** Flow cytometry plots of M. smegmatis mc^2^155::pCHERRY10 walled cells, either grown in MB7H9 or LPB (0.63 M sucrose), CWD cells, grown in LPB (1.0 M sucrose) and protoplasts (Lysozyme solution) exposed to water (control) or the detergent TX-100. Plots were divided in four quadrants representing Highly complex, non-fluorescent particles (Q1), Highly complex fluorescent particles (Q2, Walled cells), Less complex, fluorescent particles (Q3, CWD cells), and Less complex, non-fluorescent particles (Q4). Relative population percentages represented by the populations “Walled fluorescent” and “CWD fluorescent”. **B)** Quantification of flow cytometry data. White bars represent walled cells, while light grey and dark grey represent non-fluorescent particles and CWD cells, respectively. Please note that the addition of TX-100 (T), in contrast to the water control (C), dramatically reduces the number of CWD cells grown in LPB (1.0 M sucrose) and protoplasts (Lysozyme solution), while having no effect on the walled cells grown in MB7H9 and LPB (0.63 M sucrose). **C)** Quantification of Colony Forming Units (CFU) counts of walled cells (MB7H9 or LPB 0.63 M sucrose), CWD cells (LPB 1.0 M sucrose) and protoplasts (Lysozyme solution) after the exposure to TX-100 (T) or water (C). Colonies were counted after 3 days. All measurements were performed in biological triplicates and technical duplicates.

**Figure S7.**
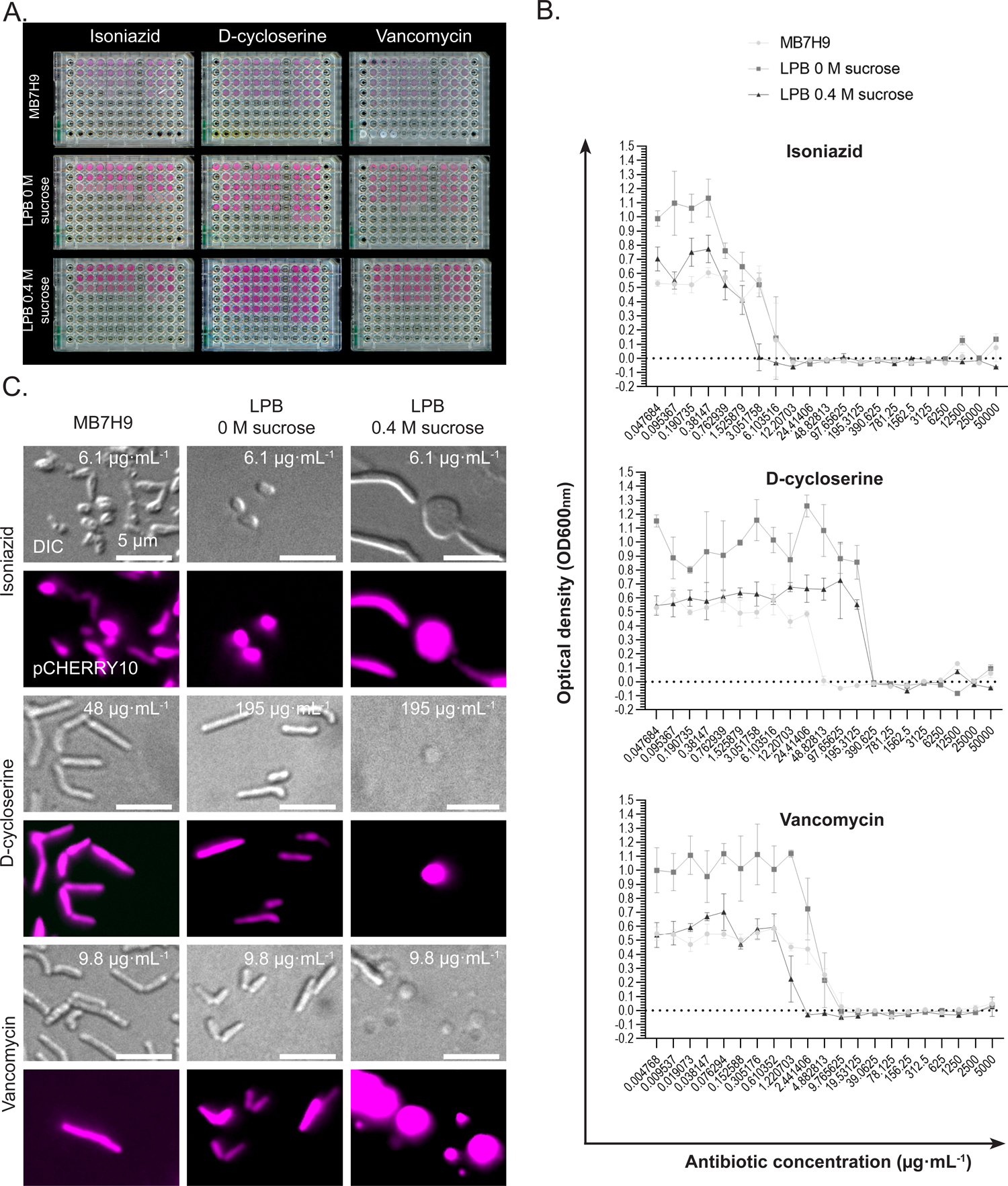
CWD formation in MIC assays of cell wall targeting antibiotics. MIC assays of *M. smegmatis* mc^2^155::pCHERRY10 in response to isoniazid, D-cycloserine and vancomyin in either MB7H9, LPB 0 M sucrose or LPB 0.4 M sucrose, performed in 96-well plates. Biological replicates were applied. Data was collected after 7 days. MIC determinations in hyperosmotic media are listed in Supplemental Table 5. **A)** Scanning images of the 96-well plates. **B)** Optical density measurements of the MIC plates. **C)** DIC-fluorescent images of walled and CWD cells. Please note that CWD cells are solely observed in sucrose supplemented LPB in response to the presence of antibiotics.

**Figure S8.**
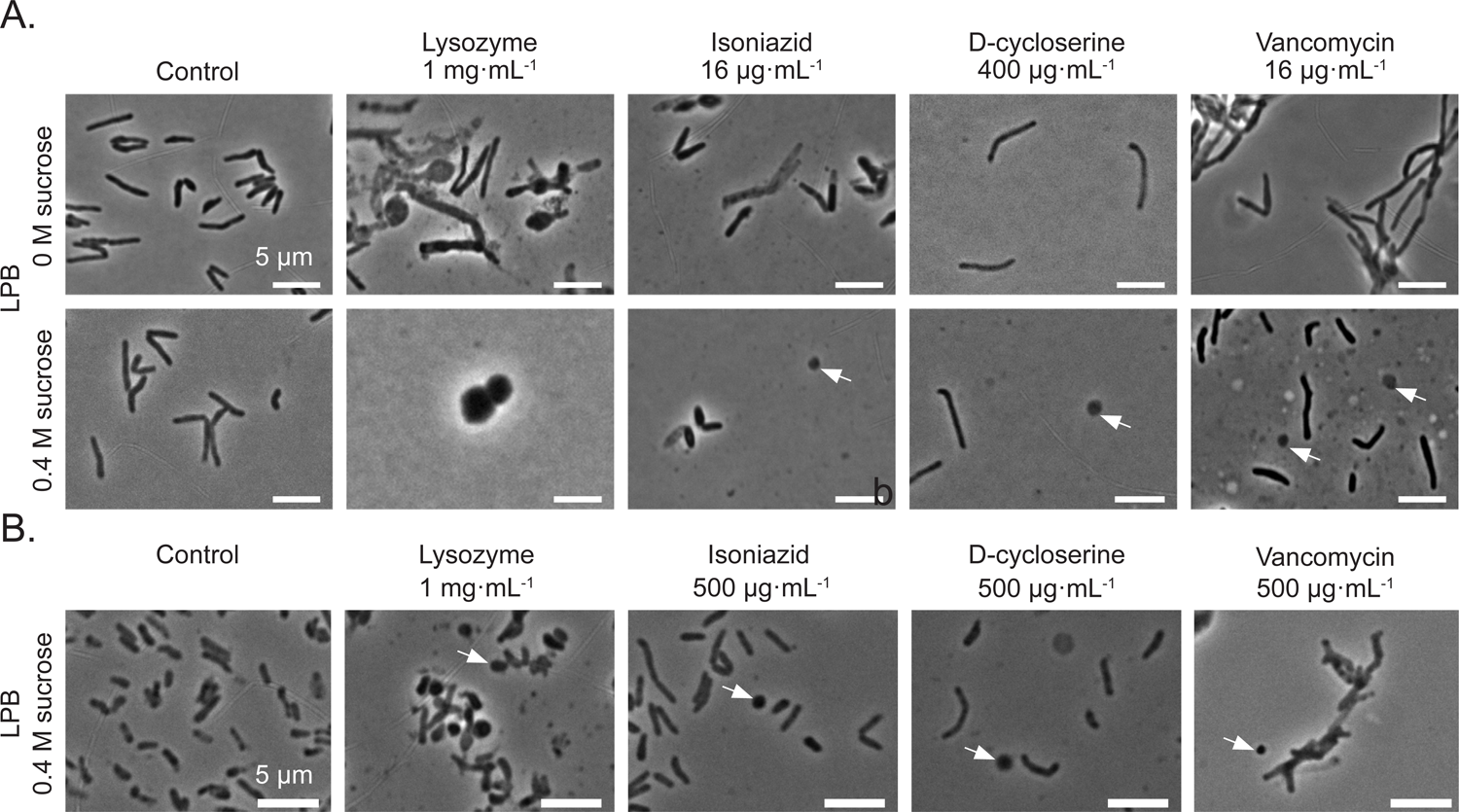
Cell wall-targeting antibiotics induces the formation of CWD cells in *M. smegmatis* Phase-contrast images showing CWD cells of *M. smegmatis* mc^2^155 formed in LPB (0.4 M sucrose) in response to cell wall-targeting antibiotics. Controls includes 1 mg/mL lysozyme + 25 mM MgCl_2_ (positive control) and no supplementation (negative control). **A)** Formation of CWD cells after 5 days exposure to either 16 µgꞏmL^-1^ isoniazid, 400 µgꞏmL^-1^ D-cycloserine or 16 µgꞏmL^-1^ vancomycin. Scale bars represent 10 µm. **B)** Formation of CWD cells after 8 days exposure to elevated antibiotic concentrations, namely 500 µgꞏmL^-1^. Scale bars represent 5 µm.

**Figure S9.**
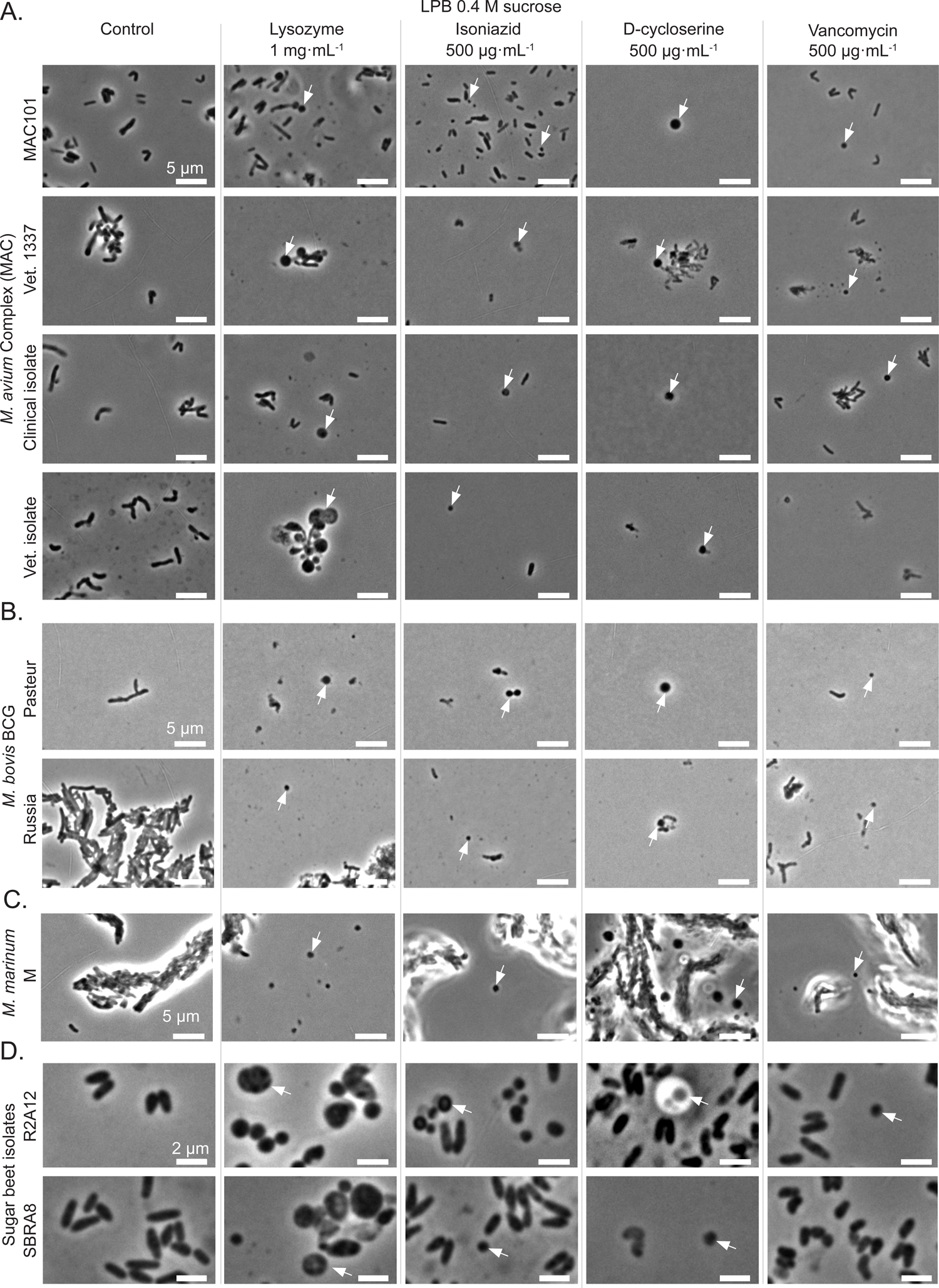
Cell wall-targeting antibiotics induces the formation of CWD cells mycobacterial species Phase-contrast images of various mycobacterial strains grown in LPB (0.4 M sucrose) supplemented with 500 µgꞏmL^-1^ of either isoniazid, D-cycloserine or vancomycin. Controls includes 1 mgꞏmL^-1^ Lysozyme + 25 mM MgCl_2_ (positive control) and no supplementation (negative Control). Scale bars represent either 5 or 2 µm. **A)** *M. avium* Complex (MAC) strains **B)** *M. bovis* BCG strains **C)** *M. marinum* M strain **D)** Endophyte isolates

**Figure S10.**
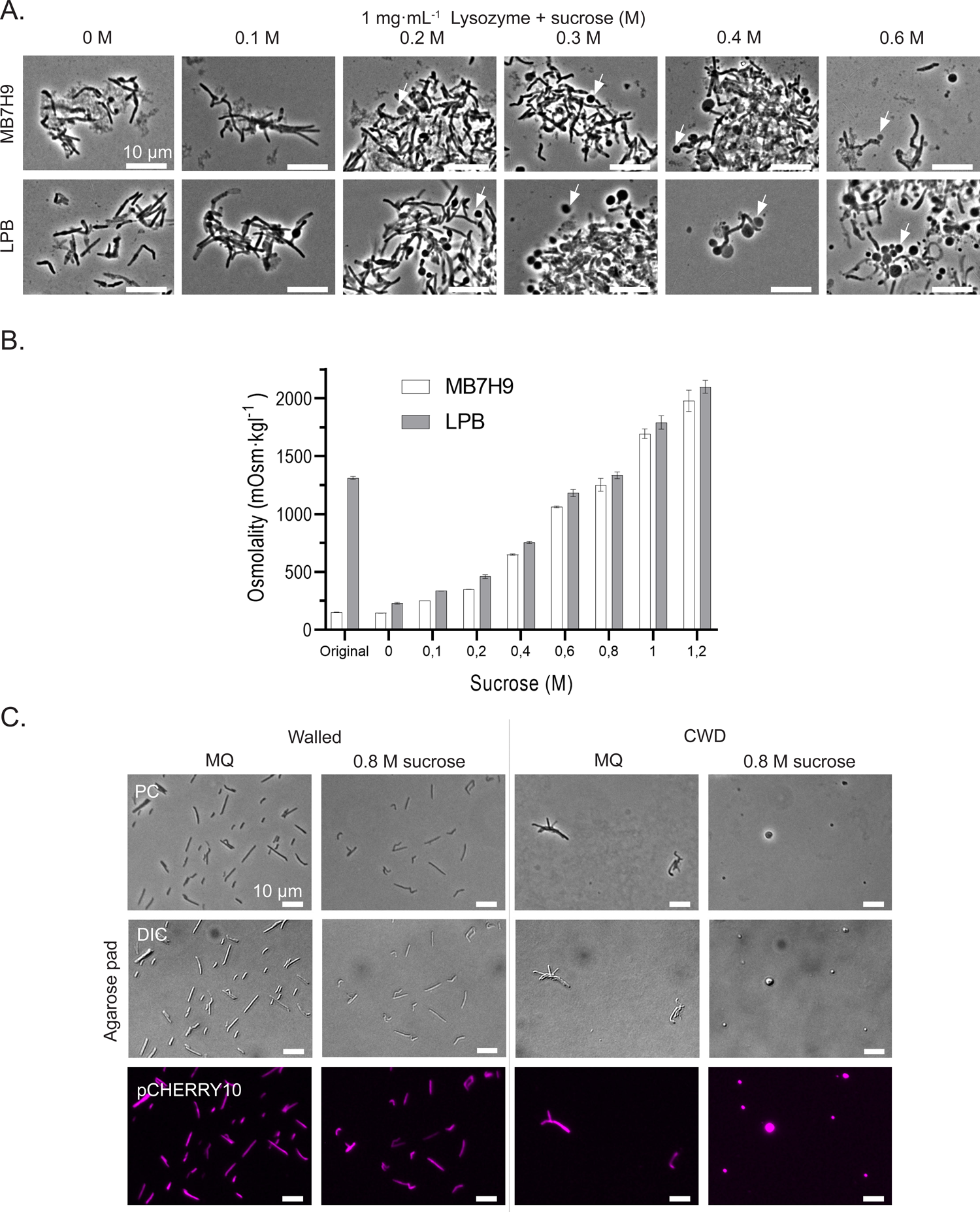
CWD cells require osmolytes. **A)** Phase contrast images of *M. smegmatis* mc^2^155 exposed to 1 mgꞏmL^-1^ Lysozyme + 25 mM MgCl2 for 24 hours in either MB7H9 or LPB supplemented with ascending concentrations of sucrose. Scale bars represent 10 µm. **B)** Osmolality measurements of LPB and MB7H9 with ascending concentrations of sucrose. Original media includes standard used LPB at RT and MB7H9 kept at 4°C. Measurements were performed in sextuplicate. **C)** Phase contrast, DIC and fluorescent micrographs of *M. smegmatis* mc^2^155::pCHERRY10 walled grown in MB7H9 and CWD cells enriched from LPB (1.0 M sucrose) mounted on MQ and MQ/sucrose (0.8 M sucrose) agar pads. Please note that all CWD cells are absent when placed on MQ agar pads. Scale bars represent 10 µm.

## SUPPLEMENTAL VIDEOS

**Video S1. CWD cell extrusion at mycobacterial polar tips** Confocal live imaging of *M. smegmatis* mc^2^155 CWD cell extrusion overtime. Exponential phase cells were grown in LPB for 6 hours prior imaging fixated under LPMA agar pad. Scale bar represent 5 µm.

**Video S2. Explosive lysis of fluorescent CWD cells in response to TX-100 treatment** Fluorescent live imaging of *M. smegmatis* mc^2^155::pCHERRY10 CWD cells in response to 0.1% TX-100 exposure. Scale bar represent 50 µm.

## SUPPLEMENTAL TABLES

**Table S1.** Mycobacterial strains used

| Strain | Genotype | Relevant description | Origin | Reference |
| --- | --- | --- | --- | --- |
| *M. smegmatis*<br>ATCC 607 | wild-type | *M. smegmatis* (Trevisan 1889) Lehmann and Neumann 1899 (DSM 43465). Human isolation source. Low transformation efficiency and strong cell clumping. | DSM | Etienne *et al.*, 2005 |
| *M. smegmatis*<br>mc<sup>2</sup>155 | ept-1 | Generally used mutated lab strain (ATCC 700084). Electro competent, loss of cell-clumping. | Wilbert Bitter | Snapper *et al.*, 1990 |
| *M. marinum*<br>M<sup>LU</sup> | wild-type | *Mycobacterium marinum* Aronson M strain (ATCC BAA-535). Isolated from Human sample (Moffet Hospital # 975973) | Annemarie Meijer | Ramakrishnan and Falkow *et al.*, 1994 |
| *M. bovis*<br>BCG<br>Pasteur | wild-type | *Mycobacterium bovis* Karlson and Lessel (ATCC 35734). Bacillus Calmette-Guérin (BCG) vaccine strain TMC 1011 [BCG Pasteur] (Pasteur 1173 P2) | ATCC | Oettinger *et al.*, 1999 |
| *M. bovis*<br>BCG Russia | wild-type | *Mycobacterium bovis* Karlson and Lessel (ATCC 35740). BCG vaccine strain TMC 1020 [BCG Russia] (Russia BCG-12) | ATCC | Oettinger *et al.*, 1999 |
| *M. avium*<br>MAC101 | wild-type | *Mycobacterium avium* Chester strain 101 (ATCC 700898). Human isolation source. | ATCC |  |
| *M. avium*<br>Vet. 1387 | wild-type | *Mycobacterium avium* supsp. *avium* Chester (ATCC 25291). Veterinary isolation strain Vet. 1387 [SSC 1336, TMC 724]. Isolated from liver of diseased hen. | Giovanni Ghielmetti |  |
| *M. avium*<br>strain<br>1904038568 | wild-type | *Mycobacterium avium*. Clinical<br>human patient isolate | Alexandra<br>Aubry | unpublish<br>ed |
| *M. avium*<br>Vet. Isolate<br>20-935 /2 | wild-type | *Mycobacterium avium* supsp.<br>*hominissuis*. Isolated from pig<br>liver. | Giovanni<br>Ghielmetti | unpublish<br>ed |
| Endophyte<br>MS5 | wild-type | Isolated from sugar beets. | NIOO-KNAW | Carrion *et al.*, 2019 |
| Endophyte<br>R2A12 | wild-type | Isolated from sugar beets. | NIOO-KNAW | Carrion *et al.*, 2019 |
| Endophyte<br>SBRA8 | wild-type | Isolated from sugar beets. | NIOO-KNAW | Carrion *et al.*, 2019 |

**Table S2.** Imaging settings used with Zeiss LSM 900 confocal microscope.

| Fluorescent protein,<br>probe or metabolic<br>label | Excitation<br>(nm) | Emission<br>(nm) |
| --- | --- | --- |
| mCherry | 561 | 570-700 |
| SYTO 9 | 488 | 490-600 |
| SynapseRed | 488 | 600-700 |

**Table S3.** Pearson chi-square tests on flow cytometry data of *M. smegmatis* grown under different conditions and exposed to TX-100. Tests are performed on averages of biological triplicates and technical duplicates. Separate populations: df = 2 Total populations: df = 4, *p-*values less than 0.05 are considered significant

|  |  | Pearson chi-squared test |  |
| --- | --- | --- | --- |
| Media | Population | X<sup>2</sup> | *p*-value |
| MB7H9 | Walled (Q2) | 0.01 | 0.99 |
|  | CWD (Q3) | 0.16 | 0.92 |
|  | Rest (Q1 & 4) | 0.01 | 0.99 |
|  | Total | 0.18 | 0.99 |
| LPB<br>0.6 M sucrose | Walled (Q2) | 0.02 | 0.99 |
|  | CWD (Q3) | 0.08 | 0.96 |
|  | Rest (Q1 & 4) | 0.25 | 0.88 |
|  | Total | 0,35 | 0.98 |
| LPB<br>1.0 M sucrose | Walled (Q2) | 0.63 | 0.73 |
|  | CWD (Q3) | 25.15 | 3.45e<sup>-06</sup> |
|  | Rest (Q1 & 4) | 26.78 | 1.53e<sup>-06</sup> |
|  | Total | 52.56 | 1.05e<sup>-10</sup> |
| Lysozyme<br>solution | Walled (Q2) | 42.69 | 5.36e<sup>-10</sup> |
|  | CWD (Q3) | 26.73 | 1.57e<sup>-06</sup> |
|  | Rest (Q1 & 4) | 462 | 4.7e<sup>-101</sup> |
|  | Total | 531 | 1.1e<sup>-113</sup> |

**Table S4.**
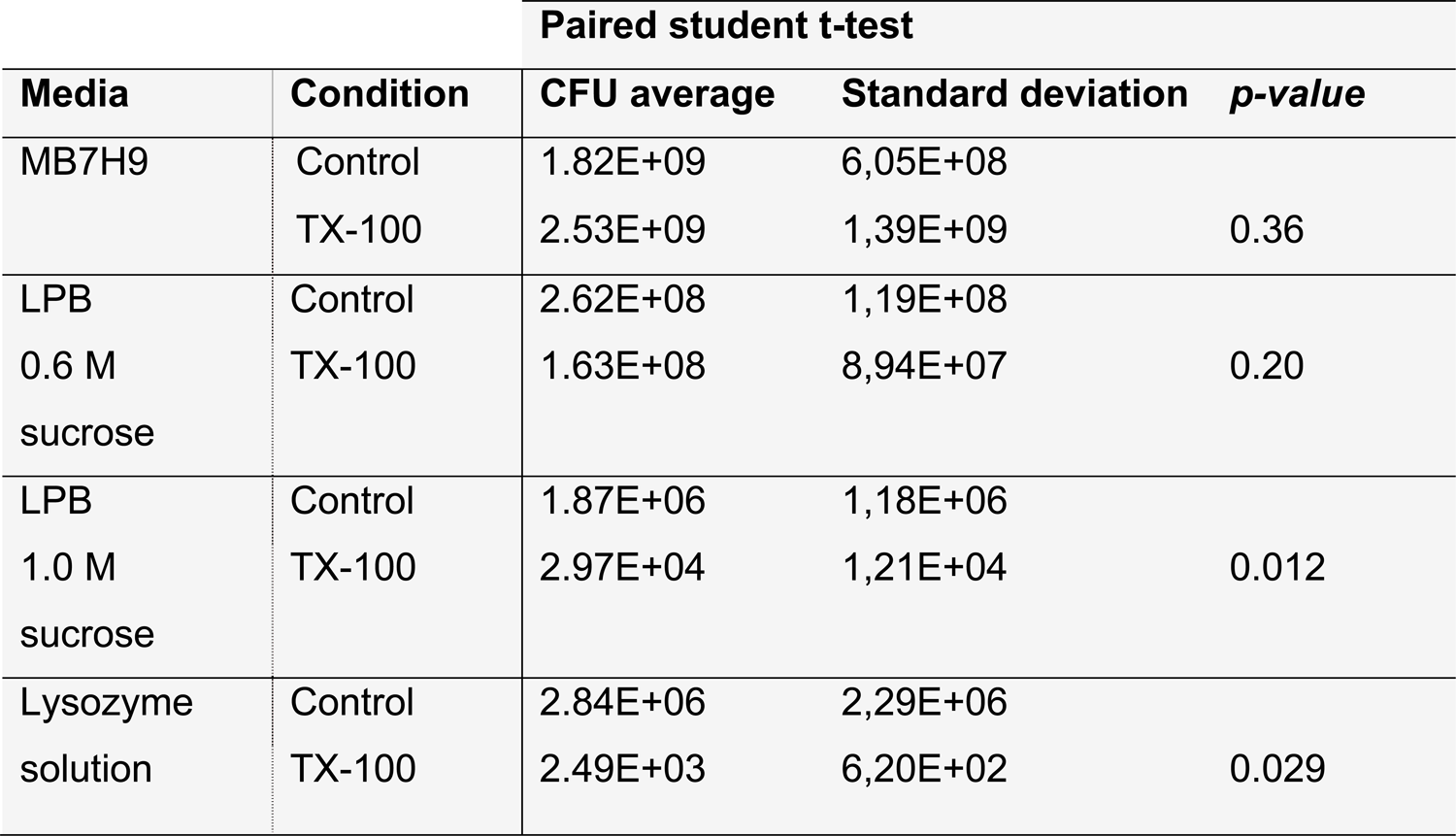
Paired student t-tests on viability assessment through CFU counting of *M. smegmatis* grown in different conditions and exposed to TX-100. Performed on biological triplicates and technical duplicates., Tails = 2, *p-*values less than 0.05 are considered significant

**Table S5.** Minimal Inhibitory Concentration (MIC) of antibiotics in hyperosmotic media

| Antibiotic | MIC ($\mu\text{g}\cdot\text{mL}^{-1}$) | | |
| --- | --- | --- | --- |
|  | MB7H9 | LPB 0 M<br>sucrose | LPB 0.4 M<br>Sucrose |
| Isoniazid | 12.2 | 12.2 | 6.1 |
| Vancomycin | 19.5 | 9.7 | 2.4 |
| D-cycloserine | 97.7 | 390.6 | 390.6 |

**Table S6.**
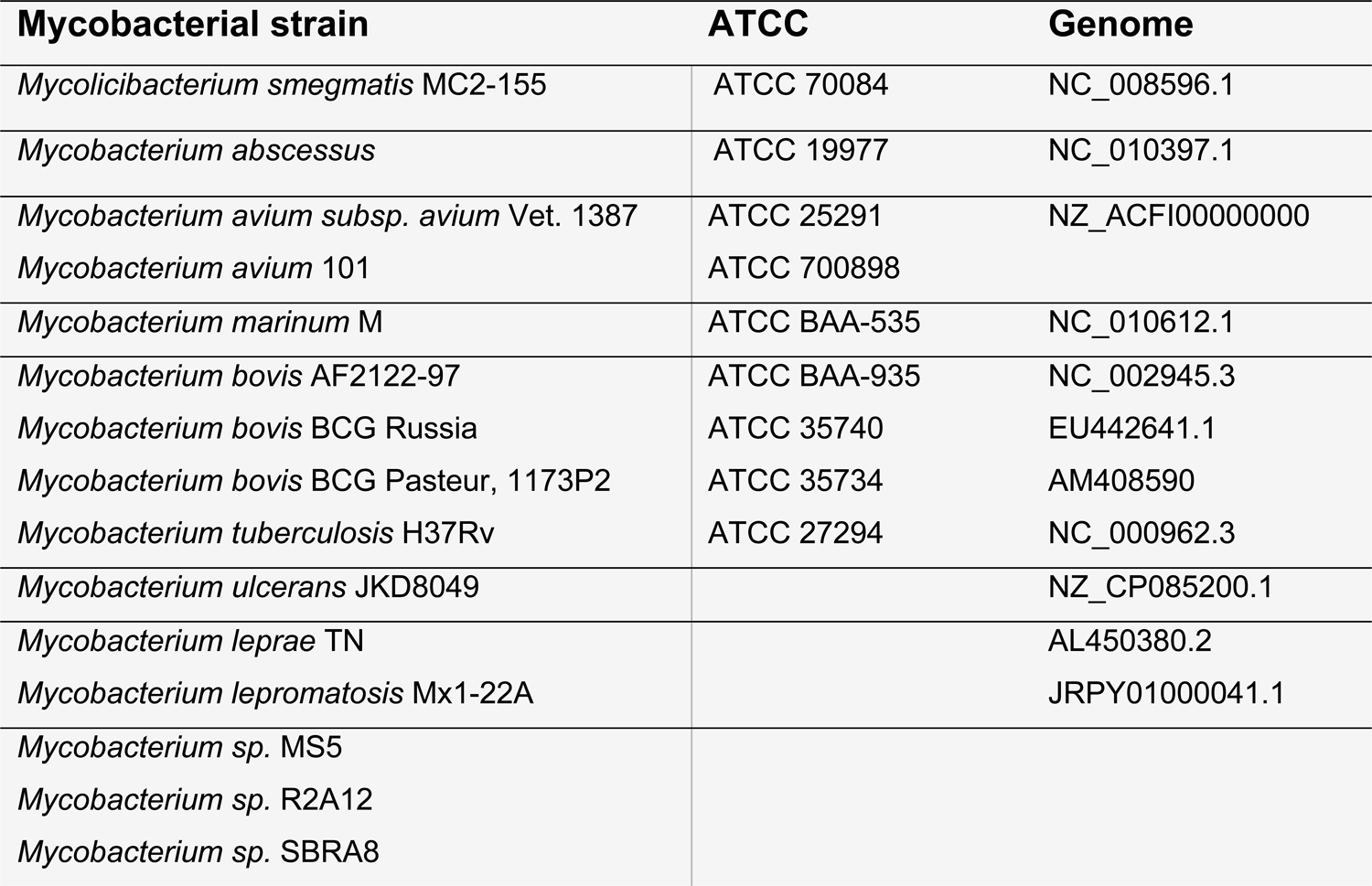
Mycobacterial genome references used for phylogenetic analysis.

**Table S7.** Osmolality measurements and osmotic pressure calculations on cultivation media supplemented with increasing concentrations of sucrose. Δosm = osmolal gap between measured osmolality and calculated osmolarity, π = osmotic pressure

| Medium | Sucrose (M) | Osmolality<br>(mOsm·kg<sup>-1</sup>) | $\Delta\text{osm}$<br>(mOsm·kg<sup>-1</sup>) | $\pi$ (mmHg) |
| --- | --- | --- | --- | --- |
| MB7H9 * | Original | 151.0 ± 1.3 | - | 2914 ± 24 |
| LPB ** | Original | 1282.5 ± 8.3 | - | 24752 ± 160 |
| MB7H9 ** | 0 | 144.7 ± 1.0 | - | 2792 ± 20 |
|  | 0.1 | 249.8 ± 0.4 | 5.2 | 4822 ± 8 |
|  | 0.2 | 348.3 ± 1.2 | 3.7 | 6723 ± 23 |
|  | 0.4 | 649.8 ± 5.1 | 105.2 | 12542 ± 99 |
|  | 0.6 | 1062.2 ± 8.4 | 317.5 | 20500 ± 162 |
|  | 0.8 | 1252.3 ± 55.5 | 307.7 | 24170 ± 1071 |
|  | 1.0 | 1693.7 ± 41.3 | 549.0 | 32688 ± 798 |
|  | 1.2 | 1979.0 ± 92.1 | 634.3 | 38195 ± 1778 |
| LPB ** | 0 | 229.8 ± 7.5 | - | 4435 ± 145 |
|  | 0.1 | 336.0 ± 8.9 | 6.2 | 6485 ± 17 |
|  | 0.2 | 459.5 ± 16.4 | 29.7 | 8868 ± 317 |
|  | 0.4 | 754.2 ± 9.7 | 124.3 | 14555 ± 187 |
|  | 0.6 | 1182.5 ± 29.6 | 352.7 | 22822 ± 571 |
|  | 0.8 | 1335.0 ± 29.7 | 305.2 | 25766 ± 574 |
|  | 1.0 | 1790.7 ± 58.1 | 560.8 | 34560 ± 1122 |
|  | 1.2 | 2099.7 ± 55.7 | 669.8 | 40524 ± 1075 |

\* Kept 4 °C, \*\* Kept RT

